# A balance between actin and Eps8/IRSp53 utilization in branched versus linear actin networks determines tunneling nanotube formation

**DOI:** 10.1101/2022.08.24.504515

**Authors:** J. Michael Henderson, Nina Ljubojevic, Thibault Chaze, Daryl Castaneda, Aude Battistella, Quentin Giai Gianetto, Mariette Matondo, Stéphanie Descroix, Patricia Bassereau, Chiara Zurzolo

## Abstract

Tunneling nanotubes (TNTs) connect distant cells and mediate cargo transfer for intercellular communication in physiological and pathological contexts. How cells generate these actin-mediated protrusions to span lengths beyond those attainable by canonical filopodia remains unknown. Through a combination of micropatterning, microscopy and optical tweezer-based approaches, we demonstrate that TNTs forming through the outward extension of actin (not through cellular dislodgement) achieve distances greater than the mean length of filopodia, and that branched Arp2/3-dependent pathways attenuate the extent to which actin polymerizes in nanotubes, limiting TNT occurrence. Proteomic analysis using Epidermal growth factor receptor kinase substrate 8 (Eps8) as a positive effector of TNTs showed that upon Arp2/3 inhibition, proteins enhancing filament turnover and depolymerization were reduced and Eps8 instead exhibited heightened interactions with the inverted Bin/Amphiphysin/Rvs (I-BAR) domain protein IRSp53 that provides a direct connection with linear actin polymerases. Our data reveals how common protrusion players (Eps8 and IRSp53) form TNTs, and that when competing pathways overutilizing such proteins and monomeric actin in Arp2/3 networks are inhibited, processes promoting linear actin growth dominate to favour TNT formation. Thus, this work reinforces a general principle for actin network control for cellular protrusions where simple shifts in the balance between processes that inhibit actin growth versus those that promote growth dictate protrusion formation and the ultimate length scales protrusions achieve.

## Main

Intercellular communication is vital for organisms, mediating across different length scales events like differentiation, tissue development/homeostasis, and wound healing. One mechanism relies on filamentous actin (F-actin)-based protrusions (*1*). Projecting a few microns from the cell, filopodia dynamically probe the environment, establishing focal adhesions or cell-cell connections for collective migration (*2*), tissue closure (*3*), and neurogenesis (*4*). Contrastingly, tunneling nanotubes (TNTs) are able to reach significantly longer distances (tens of microns) for direct communication between distant cells (*5*) and appear structurally different from filopodia (*6*).

First documented in 2004 (*7, 8*), TNTs are non-surface adherent, membranous extensions supported by F-actin that cytoplasmically connect cells allowing the transfer of cargo ranging from Ca^2+^ ions (*9*), RNAs (*10*) and plasma membrane (PM) proteins (e.g., H-Ras, Fas/CD95) (*11, 12*), to lysosomes (*13*) and mitochondria (*6, 14, 15*). This material transport ability has transformed the way we think about cell identity and the establishment of cellular networks (*16, 17*). For example, *in vivo* evidence has supported a physiological role for TNT-like structures in vascular coupling within the retina (*18*) and in facilitating cytoplasmic and membrane-bound material transfer between photoreceptors (*19, 20*), implicating nanotube-mediated maintenance in proper tissue function within the nervous system. Beyond their physiological involvement in signal transduction, apoptosis and immune responses (*12, 21*), TNTs can contribute to cancer progression (*15*), HIV-1 and SARS-CoV-2 transmission (*22, 23*), and the propagation of misfolded proteins (e.g., α-synuclein, Tau, huntingtin) involved in neurodegenerative diseases (*13, 24–26*).

For protrusions to form, F-actin remodelling processes (initiation, polymerization and bundle stabilization) need to be spatiotemporally controlled at the PM in order to work against membrane tension resistance to form a tube (*27*). Proteins like IRSp53 (insulin receptor tyrosine kinase substrate protein of 53 kDa) of the Bin/Amphiphysin/Rvs (BAR) family orchestrate membrane shape changes and actin assembly in filopodia, microvilli, and dendritic spine formation (*28*–*30*). Through its crescent-shaped inverse BAR domain IRSp53 is able to induce, sense and stabilize negative curvatures found in cellular protrusions (*31, 32*). Its CRIB (Cdc42/Rac interactive binding) domain permits regulatory control by Rho GTPases (*28, 33*), while its SH3 (Src homology 3) domain mediates interactions with actin polymerases like VASP (vasodilator-stimulated phosphoprotein) (*28, 34*) and the formin mDia1 (*35*), and nucleating promoting factors of the actin-related protein 2/3 (Arp2/3) complex like N-WASP (*36*) and WAVE2 (*35, 37*). Thus, IRSp53 self-assembly at the PM is sufficient to generate forces for protrusion growth by recruiting polymerases and locally polymerizing F-actin at IRSp53 clusters (*28, 38*).

A key partner of I-BAR proteins in filopodia and microvilli formation is Eps8 (epidermal growth factor receptor kinase substrate 8) (*33, 39, 40*). Eps8 was identified in a complex with Abi-1 (Abelson interactor 1) and Sos-1 (Son of sevenless homolog 1) that activates Rac (*41, 42*) for branched actin formation by Arp2/3 (*43*). Eps8 displays actin regulatory behaviour that depends on its binding partners: interaction with IRSp53 enhances Eps8’s F-actin bundling activity, while binding with Abi-1 prompts its capping activity (*33, 34, 44, 45*). We recently demonstrated that Eps8’s actin bundling activity promotes the formation of functional TNTs in neuronal cells (*46*), suggesting a switch in common actin-regulating components in generating shorter canonical filopodia versus longer TNTs. The mechanism(s) by which the cell uses a common pool of actin-interfacing proteins to establish length scales characteristic of TNTs is unclear.

To address this issue, we used micropatterning to control the distances over which actin-dependent processes, and not mechanisms based on cell dislodgement, lead to TNT formation at lengths beyond those of canonical filopodia. We found that Arp2/3-dependent pathways deter TNTs and that upon Arp2/3 inhibition, F-actin polymerizes over longer distances in nanotubes. Arp2/3 inhibition further promoted a specific interaction between Eps8 and IRSp53, and not with other I-BAR proteins, and their co-expression facilitated functional TNT formation. Proteomic analysis revealed that Eps8’s interactions with branched actin networks and proteins influencing filament turnover and depolymerization are reduced upon Arp2/3 inhibition, while IRSp53’s interactome remained unchanged and provided connections to needed polymerases. Our data show that protrusion length results from a competing balance between the utilization of monomeric actin and the availability of actin regulatory proteins in order to build distinct F-actin networks in a common cytoplasm.

## Results

### TNTs connect micropatterned cells at greater distances compared to filopodia

To assess TNT formation as a function of intercellular distances, we fabricated hexagonal arrays of fibronectin (FN) micropatterns to tune the separation distance *D* (Fig. 1a, b and Supplementary Fig. 1a) to be beyond the reach of filopodia which typically ranges 1–2 μm and rarely exceeds 10 μm (*1, 27*). CAD cell adherence was optimized on appropriately sized patterns to prevent unwanted cell infiltration into the surrounding PEG region (Supplementary Fig. 1b–e and Supplementary Video 1), ensuring that the connections we observe come from actin-driven mechanisms, and not from cells that migrate apart forming TNTs through cell dislodgement (*5, 47*). Overnight cell tracking (18 hr) confirmed that CAD cells remained resident to their initial micropatterns (Supplementary Fig. 1f, g, Example 1 and Supplementary Video 2), even during mitosis (Supplementary Fig. 1g, Example 3 and Supplementary Video 3), and cells rarely exhibited events of inter-micropattern translocation (Supplementary Fig. 1g, Example 2 and Supplementary Video 4).

**Fig. 1:**
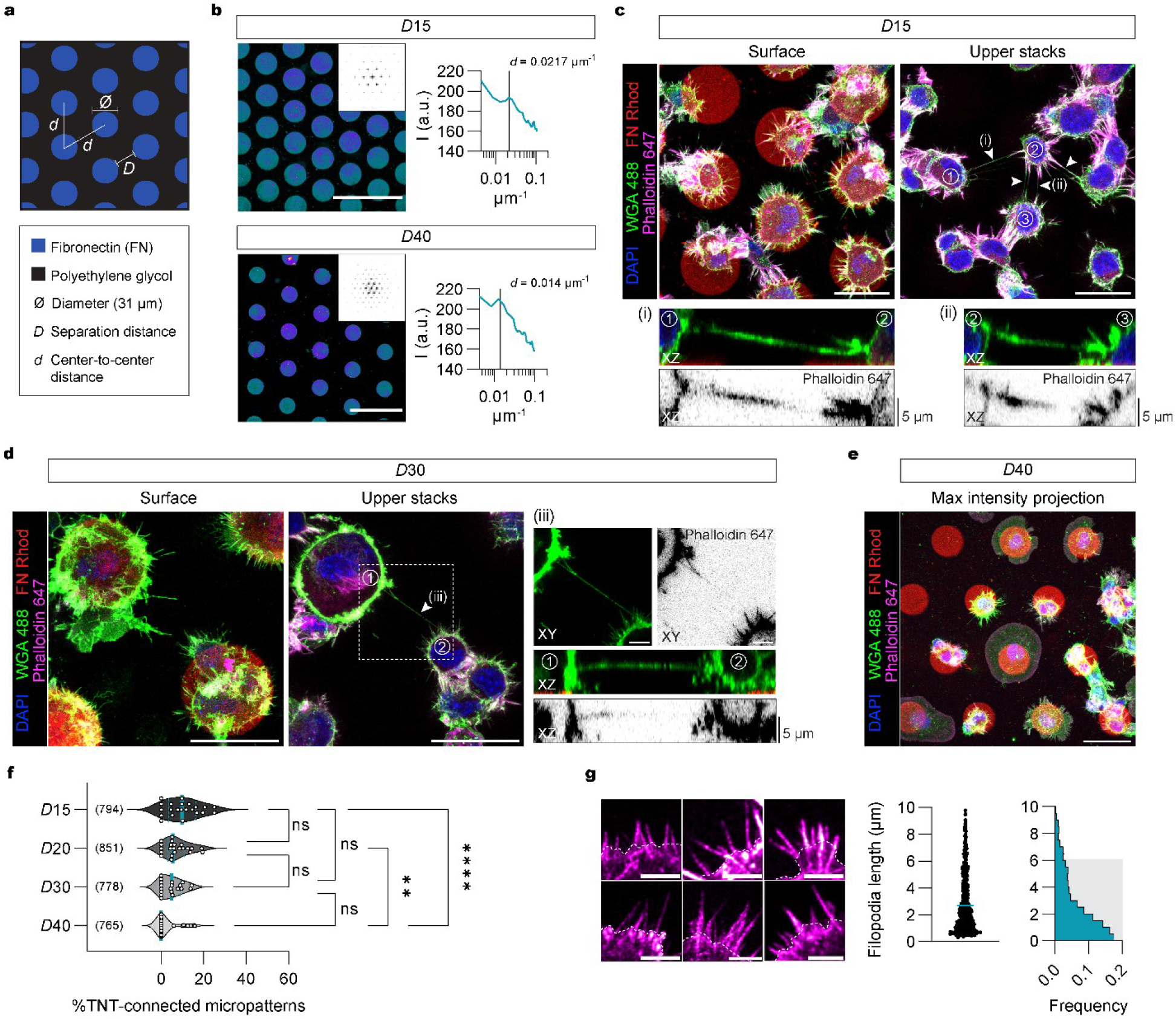
TNTs connect neighbouring CAD cells across varying distances. **(a)** Cartoon depicting fibronectin (FN) patterns produced as a hexagonal array of circles (diameter of 31 μm) with edge-to-edge separation distances (*D*) of 15, 20, 30 and 40 μm were tuned along the center-to-center distance (*d*) between circles. **(b)** Left: Representative fluorescent images of FN micropatterns spaced by 15 and 40 μm with their respective calculated fast Fourier transform (FFT) images (upper right insets). Right: Corresponding radial profiles determined from the FFT images plotted as a function of the spatial frequency (μm^−1^). Solid lines in the radial profile plots indicate the expected center-to-center *d* spacing between circular patterns and line up nicely with peaks in the measured profile, indicating high fidelity in the manufacturing process. **(c, d)** Representative surface and upper stack maximum intensity projections of CAD cells plated on 15 and 30 μm FN micropatterns. TNTs connecting cells on two different micropatterns are annotated with a white arrowhead. Subpanels (i, ii, iii) show zoom-ins and XZ projections through the axis of the indicated TNTs. **(e)** Representative maximum intensity projection of CAD cells plated on 40 μm FN micropatterns showing no TNTs between cells. **(f)** Violin plot of the percentage of TNT-connected micropatterns for *D*15, *D*20, *D*30 and *D*40 micropatterns. The median percentage of TNT-connected micropatterns (teal solid lines in the plot) on *D*15, *D*20, *D*30 and *D*40 micropatterns is 9.8%, 5.5%, 4.8%, and 0%, respectively; quartiles are annotated with dotted teal lines. Each data point corresponds to a quantified image in which on average approximately 30 cell-occupied micropatterns were within the acquired field of view; data was pooled from three experiments. The total number of individual micropatterns quantified per condition is indicated in parentheses. Statistical analysis was performed using a Kruskal Wallis test with Dunn’s multiple comparison test. ns = non-significant for P > 0.9999; **, P = 0.0074; ****, P < 0.0001. **(g)** Left: Representative images of CAD cell filopodia (magenta, Phalloidin-Alexa Flour 647). Cell body edges are marked with a dotted white line. Right: Dot plot of individual filopodia lengths (18 cells, *n* = 546 filopodia) and the corresponding histogram showing that 90% of the population (shaded gray region) had lengths that were less than 6 μm. Scale bars: (**b**) 100 μm; (**c - e**) 30 μm, Subpanels, 5 μm; (**g**) 5 μm.

We identified thin, substrate-detached TNT-like protrusions positive for F-actin connecting cells on neighbouring patterns at 15 to 30 μm (Fig. 1c, d), and containing DiD-labelled vesicles (Supplementary Fig. 2a). Using a donor-acceptor co-culture (Supplementary Fig. 2b) we found DiD vesicles both in the histone 2B (H2B)-positive acceptor cells (Supplementary Fig. 2c, Example 1) and in the connecting TNTs (Supplementary Fig. 2c, Example 2) supporting cargo transport. After increasing *D* to 40 μm, we observed fewer TNTs (Fig. 1e). *D*15 and *D*20 patterns showed the highest TNT frequency, while approaching *D*40, there was a significant decrease (Fig. 1f), indicating a distance dependency. On the other hand, filopodia lengths showed that a majority of filopodia (90% of the population) were far shorter and did not exceed 6 μm with an average length of 2.7 μm (Fig. 1g). These data show that TNTs are able to reach considerably longer lengths than average filopodia, yet there seems to be an upper limit to F-actin based elongation.

**Fig. 2:**
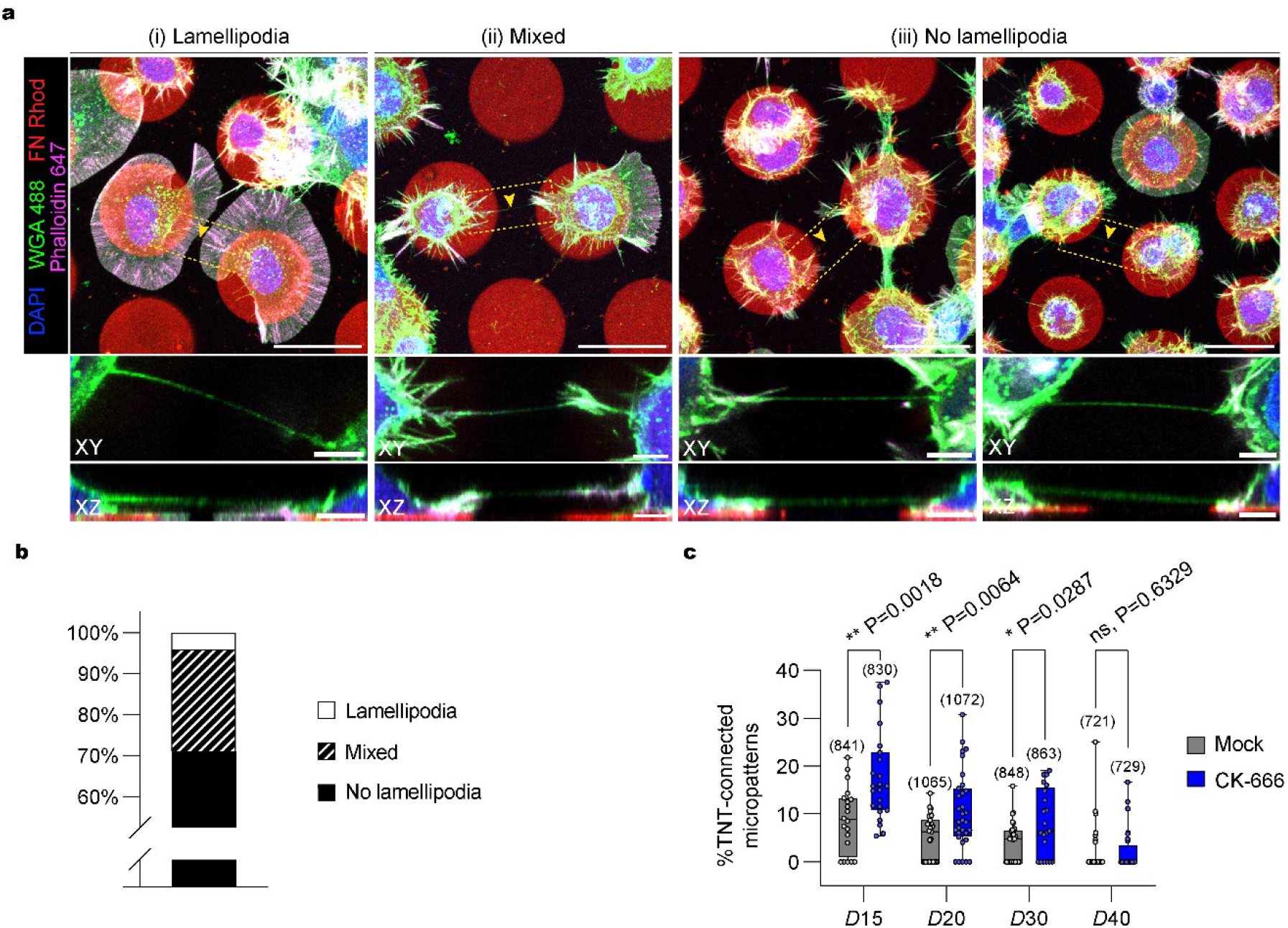
TNT prevalence negatively correlates with Arp2/3 activity. (**a, b**) TNT-connected cells characteristically lack lamellipodial features. (**a**) Representative maximum intensity projection images of cells with different morphologies that are connected by TNTs. TNTs can be formed (i) between two cells forming only lamellipodia, (ii) between two cells showing a mix of phenotypes (e.g., one of the connected cells displays only (or partial) lamellipodia while the other connected cell shows no lamellipodia), or (iii) between two non-lamellipodia cells displaying a higher density of peripheral protrusions. Dashed yellow boxes correspond to subpanels below each image that show zoom-ins and XZ projections of TNTs that were made through the axis of the connection. Yellow arrowheads directly point to the TNT. (**b**) Bar graph showing the characterization of TNTs between cells of different morphologies plated on *D*15 micropatterns (*n* = 97 TNTs). Lamellipodia, 4.1%; Mixed, 24.7%; and No Lamellipodia, 71.1%. (**c**) Whisker box plot showing the percentage of TNT-connected micropatterns for *D*15, *D*20, *D*30 and *D*40 micropatterns in cells treated with 50 μM CK-666 as compared to the mock condition (DMSO vehicle control). Mock vs. CK-666 average values were 8.8 ± 1.5% vs. 17.6 ± 1.9% for *D*15 micropatterns, 5.0 ± 0.8% vs. 10.6 ± 1.4% for *D2*0 micropatterns, 4.0 ± 0.8% vs. 8.3 ± 1.5% for *D*30 micropatterns, and 2.0 ± 0.9% vs. 2.3 ± 0.8% for *D*40 micropatterns (mean ± SEM). Each data point corresponds to a quantified image in which on average approximately 30 cell-occupied micropatterns were within the acquired field of view; data was pooled from three experiments. The total number of individual micropatterns quantified in each condition is indicated above each box. The data was analysed using an unpaired Mann-Whitney test. Scale bars, 30 μm; Subpanels, 5 μm.

### Inhibition of branched actin networks favours TNTs

Actin assembly can be organized into distinct architectures such as Arp2/3-dependent branched filaments in lamellipodia, or bundles of un-branched linear filaments. Phenotypic characterization of TNT-connected cells on *D*15 micropatterns revealed that cells displaying only lamellipodia accounted for 4.1% of TNTs examined (Fig. 2a, b). In contrast, in 71.1% of cases both cells involved in the connection had no lamellipodia (Fig 2a, b) while 24.7% of these TNT-connected cells had a mix of cellular phenotypes (Fig. 2a, b). Further analysis revealed that TNTs rarely emerge directly from lamellipodia (< 2% of cases) and instead originate either further up on the cell body or from protrusion-rich zones (Supplementary Fig. 3). Together, this suggests an anti-correlation between regions with high Arp2/3 activity and a cell’s ability to form TNTs.

**Fig. 3:**
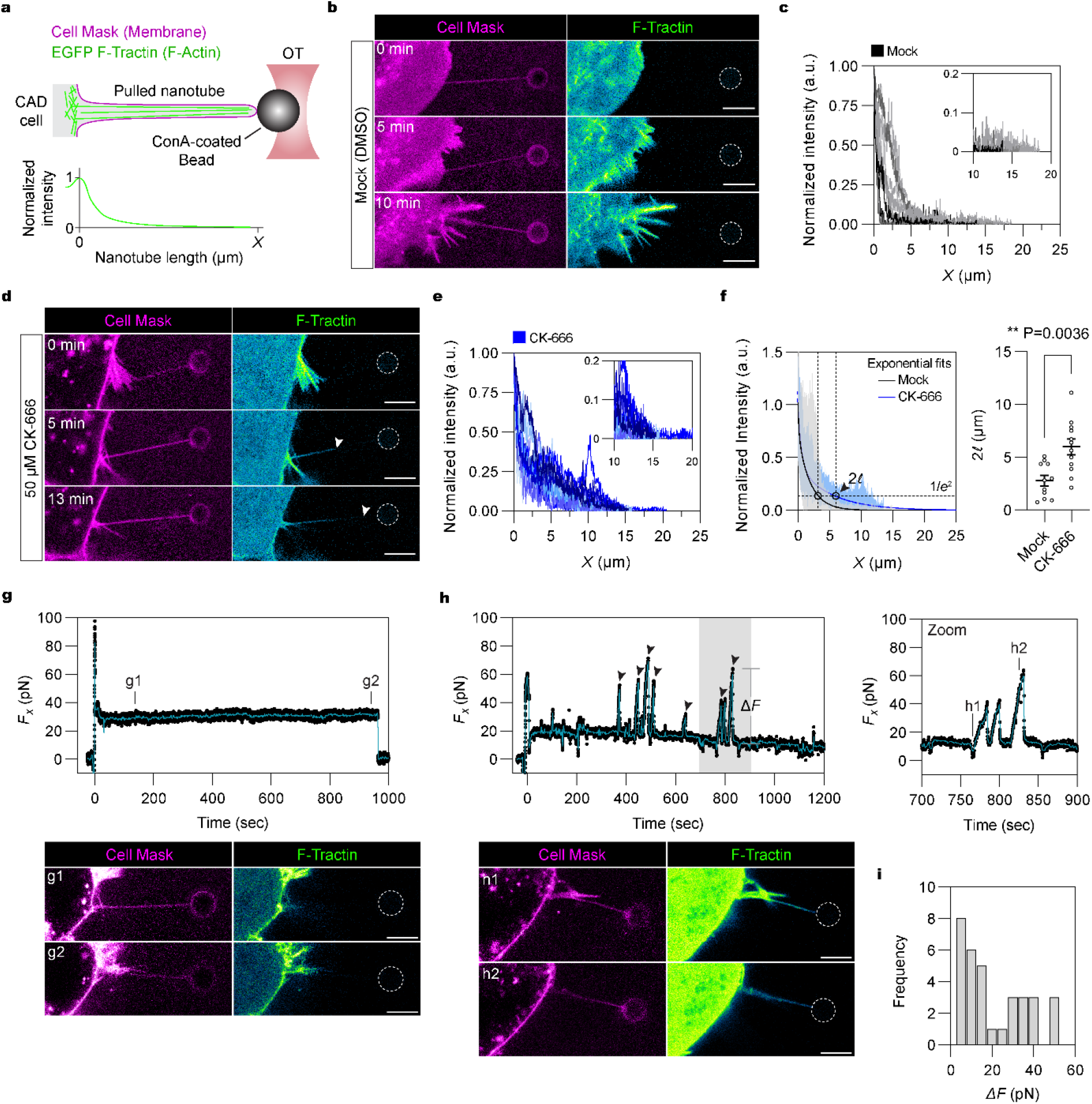
Inhibition of branched actin formation results in greater F-actin development in pulled nanotubes. (**a**) Experimental setup for pulling membrane nanotubes using a concanavalin A (Con-A)-coated bead (∼3 μm in diameter) trapped in an optical tweezer (OT). CAD cells stably expressing the small F-actin-binding peptide F-tractin fused to EGFP (green) were exogenously labelled with the lipophilic Cell Mask™ Deep Red plasma membrane stain (magenta). Intensity profiles of the F-actin fluorescence were measured along the nanotube axis and plotted against the nanotube length. (**b, d**) Representative time-lapse images of a nanotube pulled from a DMSO-treated (mock control) (Supplementary Video 5) and a CK-666-treated cell (Supplementary Video 6). White arrowheads annotate the progression of actin development within the nanotube in **d**. (**c, e**) Plot of actin profiles within pulled nanotubes for mock- and CK-666-treated cells. Insets show a magnified view at the tube extremity to better highlight the greater presence of F-actin in the CK-666 condition as compared to the mock condition. Mock condition, 11 tubes; CK-666 condition, 12 tubes. (**f**) Left: Exponential fits to the actin intensity profiles were performed to determine a characteristic decay length (2*l*) at which the initial intensity at *X* = 0 decays to a value of 1/*e*^2^. For visualization purposes of the analysis, exponential fits are shown for the mean actin profiles computed from the individual plots presented in **e** and **f**. Upper and lower limits of the intensity range are shaded. Right: Dot plot of the characteristic decay lengths (2*l*) for mock- and CK-666-treated cells. Data is represented as the mean ± SEM. Mock (11 tubes), 2.78 ± 0.50; CK-666 (12 tubes), 5.80 ± 0.73. Statistical analysis was performed using an unpaired Mann-Whitney test. (**g**) Top: Force plot of a pulled nanotube from a mock-treated cell showing no F-actin development. Solid teal line, 10-point moving average curve. Bottom: Associated images of the indicated time points (g1, g2). (**h**) Top: Force plot of a pulled nanotube from a CK-666-treated cell showing F-actin development spanning the entire nanotube length. Peaks in the force plot (black arrowheads), with magnitudes of Δ*F*, arise when retrograde flows outcompete actin polymerization (at the nanotube tip) causing bead displacement towards the cell body (recorded as a positive rise in the force in the lab frame). Solid teal line, 10-point moving average curve. Shaded grey region corresponds to a magnified view on the right. Bottom: Associated images of the indicated time points (h1, h2). (**i**) Histogram of the force peak magnitudes (Δ*F*). Sample size, 33 peaks. The trapped bead is annotated by a dotted white circle when not clearly visible. Scale bars, 5 μm.

We next assessed if the inhibition of actin incorporation in Arp2/3-dependent networks favours the formation of TNTs. Upon Arp2/3 inhibition with CK-666 (*6, 48*), we observed a significant increase in the percent of TNT-connected cells on *D*15, *D*20 and *D*30 micropatterns (Fig. 2c). In contrast, CK-666 had no impact on the number of TNT-connected cells on *D*40 patterns (Fig. 2c) again suggesting there is an upper limit to the length a TNT can obtain through the polymerization of actin (Fig. 1f and Fig. 2c). We hypothesized that a competition between different actin networks has a direct effect on the likelihood for a TNT to form over a given distance. Thus we directly shifted the actin balance in favour of linear F-actin by using the formin-specific agonist drug IMM-01 (*49*) and assessed the effect on TNTs. With IMM-01 treatment, we observed an increased number of TNT-connected cells (Supplementary Fig. 4a, b) and of vesicles transferred from donor to acceptor cells (Supplementary Fig. 4c), indicating that the promotion of linear actin networks directly favours functional TNT formation. Together this data suggests that TNT formation is based on the polymerization of long, linear filaments and becomes enhanced when branched actin formation is inhibited.

**Fig. 4:**
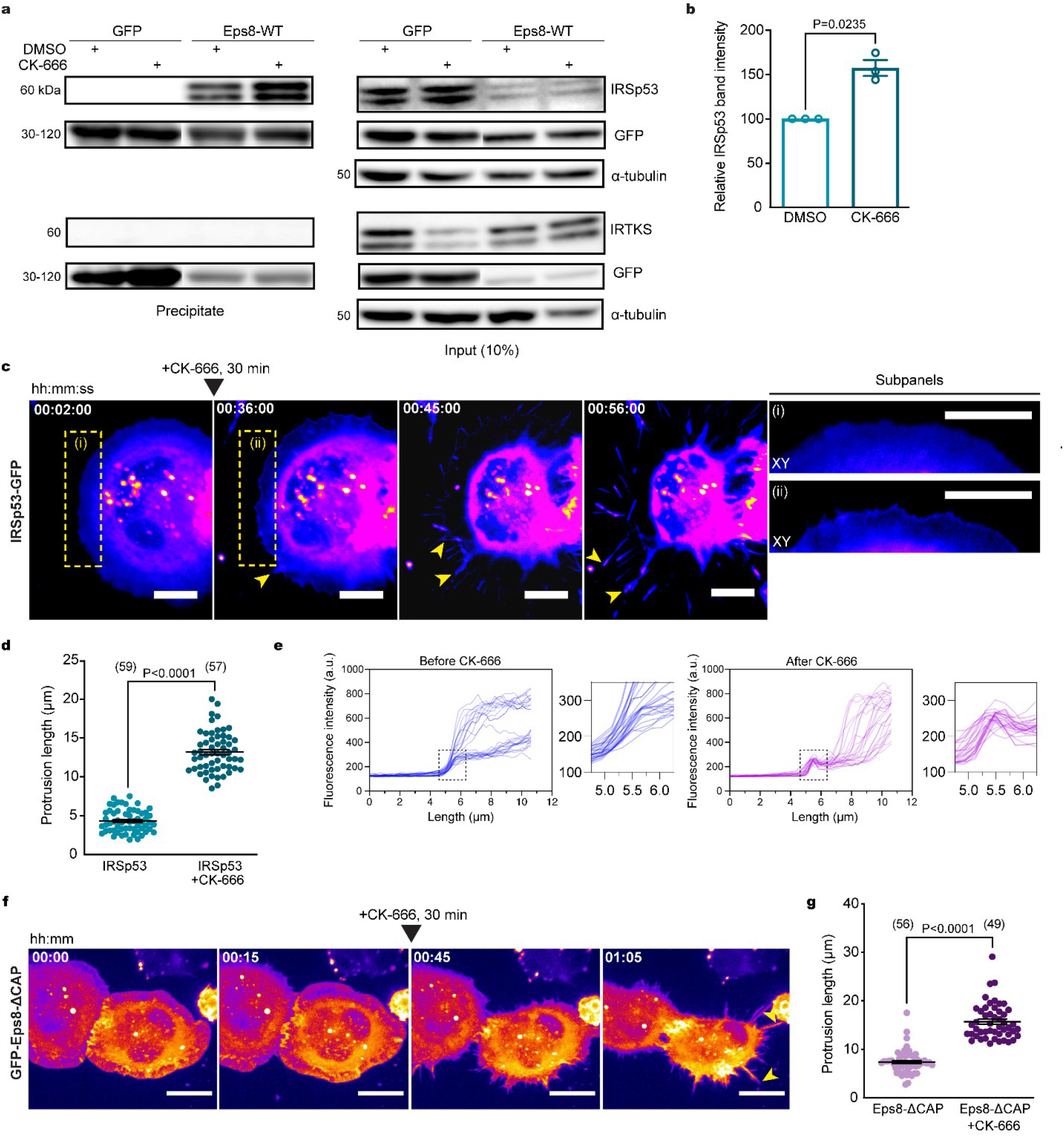
Eps8 and IRSp53 are recruited to form longer protrusions upon Arp2/3 inhibition. **(a)** Representative Western Blots showing GFP-trap immunoprecipitation bands of the IRSp53 and IRTKS pray proteins in the GFP-Eps8-WT bait precipitate (left) and input lysate (right), with or without 50 μM CK-666 treatment. **(b)** Relative IRSp53 band intensity quantification. Both the CK-666-treated and DMSO-treated bands for IRSp53 were normalized with respect to their Eps8-WT bait bands (detected by an anti-GFP antibody). Relative band intensity was 157.5 ± 8.9% for the CK-666 condition as compared to the DMSO control (was set to 100%). Data are from three individual experiments and are represented as a mean ± SEM. Statistical analysis was performed using a t-test with Welch’s correction, P = 0.0235. **(c)** Representative time-lapse images (2, 36, 45 and 56 min from Supplementary Video 7) of protrusion formation in IRSp53-transfected cells before and after CK-666 addition (at 30 min). Yellow arrowheads show protrusions that are formed after CK-666 addition. Subpanels (i, ii) show the enlarged sections of the cell edge before (i) and after (ii) the addition of CK-666, demonstrating the recruitment of IRSp53 at the membrane (ii). **(d)** Scatter plot comparing the maximum protrusion length before (4.3 ± 0.2 μm) and after CK-666 addition (13.2 ± 0.3 μm) in IRSp53-transfected cells containing the F-actin label tdTomato-F-tractin. Twenty-five cells were analysed, and the number of protrusions analysed in each condition is indicated in parentheses. Data are from 10 individual experiments and are represented as a mean ± SEM. Statistical analysis was performed using an unpaired t-test with Welch’s correction, P < 0.0001. (**e**) Profile plots of IRSp53 intensity determined across the membrane before (left) and after the addition of CK-666 (right). Magnified views of the cell edge (dashed black boxes) are provided to better highlight the recruitment of IRSp53 at the cell membrane in the CK-666 condition as compared to the control. Two cells were analysed: before CK-666, 28 measurements; after CK-666, 25 measurements. **(f)** Representative time-lapse images (0, 15, 45 and 65 min from Supplementary Video 8) of protrusion formation in Eps8-ΔCAP-transfected cells before and after CK-666 addition (at 30 min). Yellow arrowheads show protrusions that are formed after CK-666 addition. **(g)** Scatter plot comparing the maximum protrusion length before (7.4 ± 0.3 μm) and after CK-666 addition (15.7 ± 0.5 μm) in Eps8-ΔCAP-transfected cells containing the F-actin label tdTomato-F-tractin. Twenty-one cells were analysed, and the number of protrusions analysed in each condition is indicated in parentheses. Data are from 9 individual experiments and are represented as a mean ± SEM. Statistical analysis was performed using unpaired t-test with Welch’s correction and P value < 0.0001. Scale bars: (**c**) 10 μm, Insets, 10 μm; (**e**) 20 μm.

### F-actin polymerization occurs over longer distances when Arp2/3 is inhibited

To gain a greater insight into how TNTs can reach longer distances, we utilized an optical tweezer (OT) setup to pull nanotubes of comparable lengths to the TNTs observed on the micropatterns to monitor by confocal microscopy F-actin polymerization within the nanotube in control and CK-666-treated conditions (Fig. 3a) (*38, 50–52*). In the mock condition, cells displayed little to no F-actin within the nanotubes (Fig. 3b, c and Supplementary Video 5). In contrast, CK-666-treated cells showed a robust development of F-actin (Fig. 3d, e and Supplementary Video 6). Profiles of the F-actin fluorescence showed a rapid decay of the signal in mock-treated cells where intensity values were indistinguishable from background levels beyond 5 μm; however, CK-666-treated cells exhibited a shallower decay and intensities remained higher throughout the entire nanotube (Fig. 3c, e).

Exponential fitting of the actin profiles (Fig. 3f) and force measurements (Fig. 3g, h) confirmed CK-666-treated cells had F-actin reaching significantly longer distances within the nanotube compared to mock conditions. For example, force profiles remained static over time in control conditions for actin-devoid nanotubes (Fig. 3g), whereas large bead displacements (i.e., a rise in the force, Δ*F*) were observed in the CK-666 condition (Fig. 3h), indicating that F-actin development had reached the end of the tube, imparting large traction forces (Fig. 3i). This characteristic actin-dependent behaviour has been described previously by us and others (*38, 50, 51, 53*). These data indicate that an inhibition of Arp2/3 activity directly correlates with the polymerization of longer actin filaments.

### An Eps8-IRSp53 interaction is enhanced upon Arp2/3 inhibition

A shift in actin utilization upon Arp2/3 inhibition likely promotes interactions that outwardly deform the PM and polymerize F-actin locally for TNT outgrowth. Given Eps8’s previously identified TNT-promoting capability (*46*), we investigated whether a specific Eps8-I-BAR interaction favours long TNT formation. We first screened Eps8’s interaction with IRTKS (insulin receptor tyrosine kinase substrate) and IRSp53 based on their I-BAR domain sequence similarity (*32*) and their involvement in the formation of filopodia and microvilli (*28, 29, 39, 40*). GFP-Eps8-wild type (WT) was used to immunoprecipitate endogenous IRTKS and/or IRSp53 in control and TNT-promoting conditions using CK-666 (Supplementary Fig. 5a). IRSp53 precipitated with GFP-Eps8-WT in control conditions, indicating their direct interaction in CAD cells, while IRTKS did not (Fig. 4a). To see if the interaction is affected by Arp2/3 inhibition, we treated GFP-Eps8-WT cells with CK-666. A significant increase in eluted IRSp53 was observed, whereas no effect was observed for IRTKS (Fig. 4a, b). These data indicated that interaction between Eps8 and IRSp53, but not IRTKS, would favour long TNT growth.

**Fig. 5:**
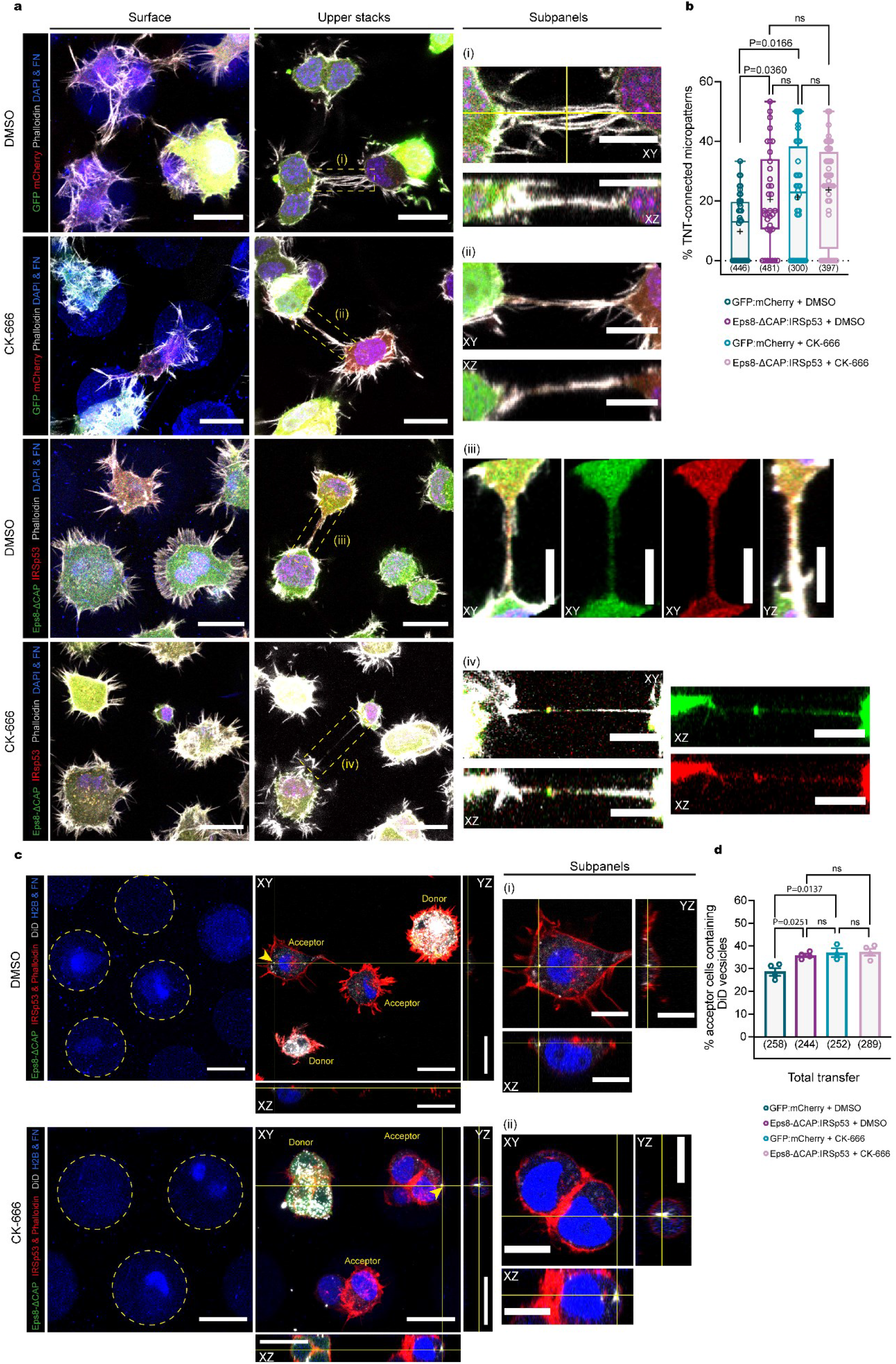
Eps8–IRSp53 co-expression leads to the formation of functional TNT-like protrusions on micropatterns. **(a)** Representative surface and upper stack images of control GFP:mCherry and GFP-Eps8-ΔCAP:IRSp53-mCherry expressing CAD cells plated on *D*15 micropatterns (Alexa 405-labelled FN) treated with either DMSO or CK-666. Subpanels (i–iv) show magnified projections of the TNTs indicated in the dashed yellow boxes; XZ and YZ projections were made through the axis of the TNT. **(b)** Whisker box plot of the percentage of TNT-connected cells on *D*15 micropatterns for GFP:mCherry + DMSO (10.3 ± 1.6%), Eps8-ΔCAP:IRSp53 + DMSO (20.8 ± 2.8%), GFP:mCherry + CK-666 (21.4 ± 3.1%), and Eps8-ΔCAP:IRSp53 + CK-666 cells (23.5 ± 2.4%) (mean ± SEM). Total number of quantified cells on patterns is indicated below the Whisker plots for each condition. Data were pooled from 3 individual experiments and each data point corresponds to a quantified image in which on average approximately 10 cells were within the acquired field of view. Mean values are indicated as a + symbol on the graph for each condition. Statistical analysis was performed using a Kruskal Wallis test with Dunn’s multiple comparison test. Significant P values for each comparison are stated on the plot, ns = non-significant for P > 0.9999. **(c)** Representative images of GFP-Eps8-ΔCAP:IRSp53-mCherry donor cells co-cultured with H2B-EBFP acceptors treated with either DMSO or CK-666. Dashed yellow circles annotate Alexa 405-labelled FN patterns for clarity and yellow arrowheads show the donor-originating vesicle in the acceptor cell. Subpanels (i, ii) show magnified projections of the acceptor cell containing DiD-labeled vesicles (yellow arrowheads); XZ and YZ projections were made through the vesicle. **(d)** Bar graph showing the percentage of H2B-EBFP acceptors containing DiD-stained vesicles in GFP:mCherry + DMSO (28.8 ± 1.7%), Eps8-ΔCAP:IRSp53 + DMSO (35.7 ± 0.7%), GFP:mCherry + CK-666 (37.0 ± 1.9%), and Eps8-ΔCAP:IRSp53+CK-666 (37.2 ± 1.5%) experiments. Total number of quantified acceptor cells is indicated for each condition. Data are from at least 3 individual experiments and are represented as a mean ± SEM. Statistical analysis was performed using an ordinary ANOVA with Tukey’s multiple comparison test. Significant P values for each comparison are stated on the bar graph. ns = non-significant for GFP:mCherry + CK-666 vs. Eps8-ΔCAP:IRSp53 + DMSO, P = 0.9246; for GFP:mCherry + CK-666 vs. Eps8-ΔCAP:IRSp53 + CK-666, P = 0.9999; and for Eps8-ΔCAP:IRSp53 + DMSO vs. Eps8-ΔCAP:IRSp53 + CK-666, P = 0.8807. Scale bars, 20 μm; Subpanels, 10 μm.

Using spinning disk microscopy, we then characterized the process of IRSp53- and Eps8-driven protrusions. IRSp53 and Eps8 in its bundling active form (capping defunct mutant; Eps8-ΔCAP) were separately introduced in CAD cells, and the maximum length of cell-generated protrusions was analysed before and after CK-666 addition (Fig. 4c–g). Prior to CK-666 treatment (first 30 min), IRSp53- and Eps8-ΔCAP-expressing cells were found to form lamellipodia devoid of protrusions (Fig. 4c, f and Supplementary Videos 7, 8) or shorter filopodia-like protrusions (Supplementary Fig. 6a, b and Supplementary Videos 9–12). After CK-666 addition, new IRSp53- and Eps8-positive protrusions grew two to three times longer than those found in the non-treated condition (Fig. 4d, g). Following CK-666 addition, we observed an increase of IRSp53 fluorescence at the plasma membrane (Fig. 4e, Supplementary Video 7) which then coincided with the generation and growth of IRSp53-positive dorsal protrusions, indicating a greater availability of IRSp53 to generate protrusions when branched actin pathways are inactivated.

**Fig. 6:**
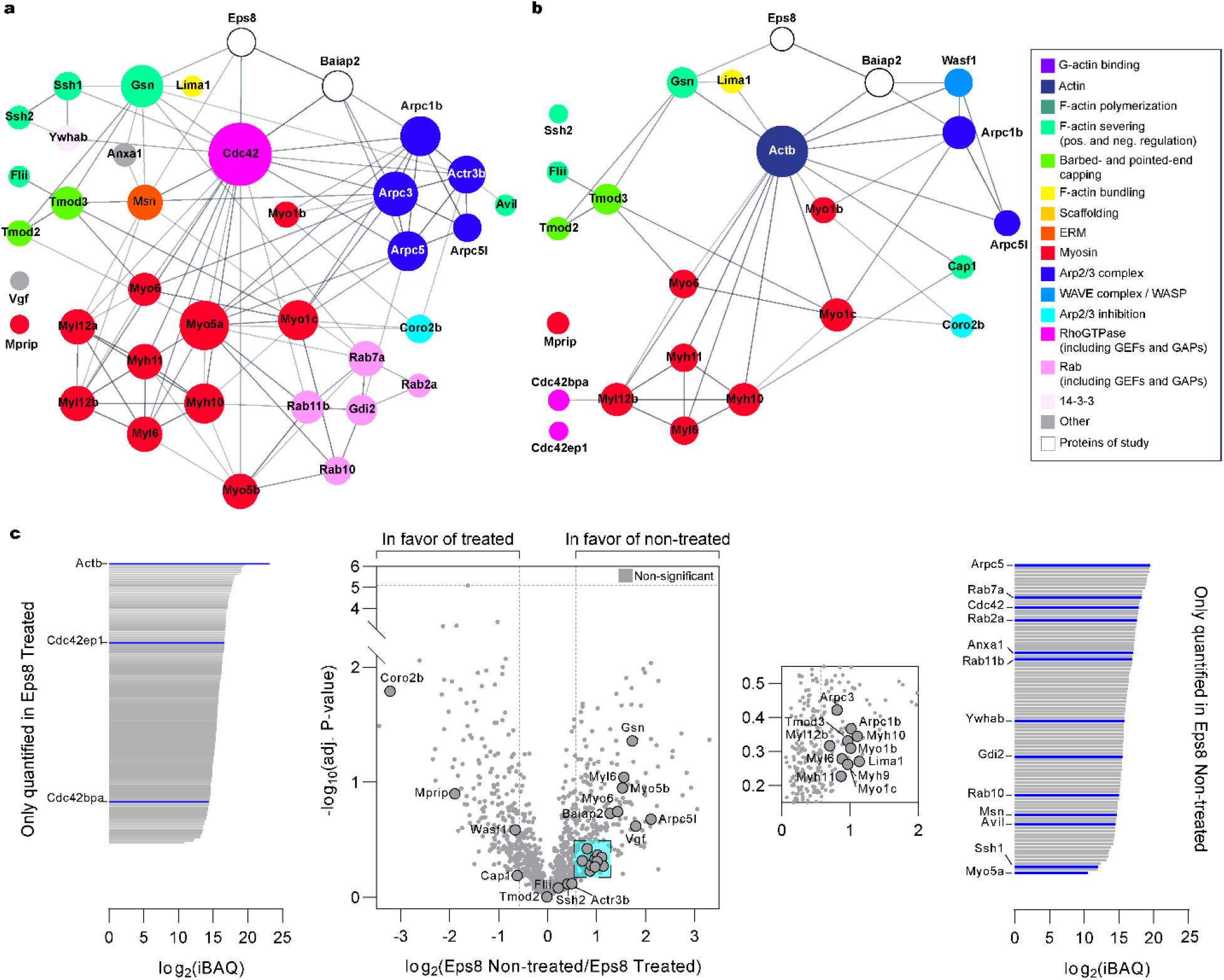
Eps8’s interactome reveals a weakened interaction with Arp2/3 complex components and proteins affecting filament turnover and depolymerization upon CK-666 treatment. **(a)** STRING-generated networks of differentially abundant proteins present in Eps8-WT as compared to the negative control for non-treated (DMSO) (**a**) and CK-666-treated (**b**) CAD cells. The sizes of the nodes (i.e., protein gene names) were scaled based on the degree of connectivity in the networks, and the transparency of the edges relate to the calculated score of the protein-protein interaction in the STRING database. (**c**) Direct comparison of identified protein hits between non-treated and treated Eps8 samples. Left: iBAQ plot of proteins only present in the Eps8 treated sample. Center: Volcano plot of common proteins to both non-treated and treated Eps8 samples. Dashed vertical lines mark the binary logarithm position of a fold change of 1.5, and the dashed horizontal lines marks the adjusted P-value threshold. Cyan-shaded square corresponds to a magnified view of the indicated region on the right of the volcano plot. Right: iBAQ plot of proteins only present in the Eps8 non-treated sample.

### Eps8 and IRSp53 co-expression increases the formation of functional TNTs

Together the above data suggest that Eps8 and IRSp53 specifically drive long protrusion formation and that their interaction is enhanced upon Arp2/3 inhibition. To further explore this hypothesis, we tested the effect of Eps8 and IRSp53 co-expression on TNT formation and their ability to transfer vesicles (donor-acceptor assay). Stable co-expression of GFP-Eps8-ΔCAP and IRSp53-mCherry led to a doubling in the percentage of TNT-connected cells on *D*15 micropatterns (20.8%) compared to the GFP:mCherry control cells (10.3%) (Fig. 5a, b). Notably, TNTs found in Eps8-ΔCAP:IRSp53 cells were positive for both proteins all along their length (Fig. 5a, inset iii). Furthermore, vesicle transfer from Eps8-ΔCAP:IRSp53 donors to EBFP-H2B acceptors also increased (Fig. 5c, d), while secretion-based transfer remained unchanged (Supplementary Fig. 7). As a control we overexpressed IRTKS (Supplementary Fig. 5b) and showed it had no positive impact on the percentage of TNT-connected cells (Supplementary Fig. 5c) nor on the transfer of vesicles (Supplementary Fig. 5d). Similarly, the co-expression of IRTKS with Eps8-WT and Eps8-ΔCAP showed a significant decrease in the percentage of TNT-connected cells (Supplementary Fig. 5e, f). Interestingly, the addition of CK-666 to Eps8-ΔCAP:IRSp53 cells had no further enhancement on the percentage of TNT-connected cells nor on vesicle transfer as compared to control GFP:mCherry cells (Fig. 5b, d), suggesting that the overexpression of Eps8 with IRSp53 may already saturate the cellular ability for TNTs to form.

**Fig. 7:**
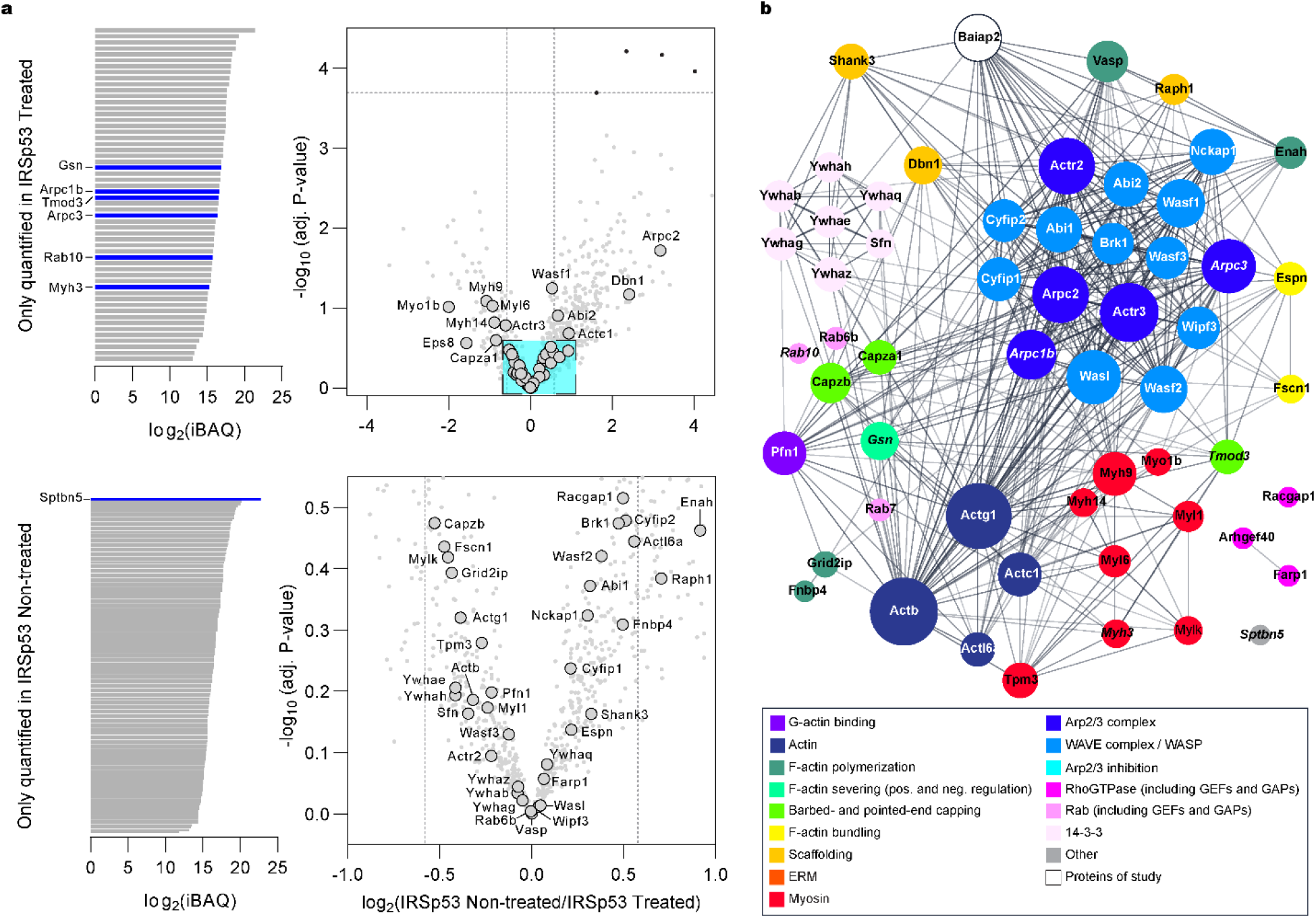
IRSp53’s interactome remains invariant to Arp2/3 inhibition. **(a)** Direct comparison between non-treated and treated IRSp53 samples. Left: iBAQ plot of proteins only present in the treated (top) and non-treated (bottom) IRSp53 samples. Right: Volcano plots of common proteins to both non-treated and treated IRSp53 samples. Dashed vertical lines mark the binary logarithm position of a fold change of 1.5, and the dashed horizontal line marks the adjusted P-value threshold. Cyan-shaded square corresponds to a magnified view of the indicated region. **(b)** Combined STRING-generated network of all identified protein hits in the IRSp53 pull down (both non-treated and treated samples). The sizes of the nodes (i.e., protein gene names) were scaled based on the degree of connectivity in the networks, and the transparency of the edges relate to the calculated score of the protein-protein interaction in the STRING database. Italicised hits are those proteins present only in the treated (e.g., Gsn, Arpc1b, Arpc3, Tmod3, Rab10, Myh3) or non-treated (e.g., Sptbn5) IRSp53 sample shown in **a**.

### Eps8 is removed from pathways associated with filament turnover upon Arp2/3 inhibition

To analyse the actin-centric pathways through which Eps8 and IRSp53 function, GFP-Eps8-WT or IRSp53-GFP were utilized as bait proteins in a whole-cell GFP-trap pull-down and the interactomes were assessed against appropriately-treated cytosolic GFP-expressing cells (negative controls) to identify protein hits (Supplementary Fig. 8a). In the Eps8-WT pull-down, we identified 36 proteins (out of 145) (Supplementary Fig. 8b), and the online Search Tool for Recurring Instances of Neighbouring Genes (STRING) database (*54*) was used to generate association networks where individual proteins were grouped into actin-based modules (Fig. 6a). As expected, in control conditions Eps8 preferentially interacts with IRSp53 (Baiap2), while no other I-BAR proteins were identified. Consistent with Eps8’s role in membrane trafficking, several Rab-related proteins were observed, e.g., Rab5 (*55*). In addition, our analysis revealed novel hits in Eps8’s interactome. We observed enrichment of Arp2/3 complex proteins (Arpc1b, Arpc3, Arpc5, Arpc5l and Actr3b). We also noted proteins directly affecting F-actin dynamics and organization, including: (1) severing proteins (gelsolin (Gsn), advillin (Avil) and flightless 1 (Flii) (*56*)) and phosphatases (Slingshot homology 1 and 2 (Ssh1 and Ssh2)) that activate cofilin (*57*); (2) pointed-end capping proteins (tropomodulins 2 and 3 (Tmod2 and Tmod3)) (*58*); (3) the bundling protein eplin (Lima 1) (*59*); and (4) coronin 2b (Coro2b) that disassembles actin branches (*60*). Finally, we observed significant pull-down of myosin-II complex proteins, consisting of both light (Myl6, Myl12a/b) and heavy (Myh10, Myh11) chain proteins, unconventional myosin motors (Myo5a/b, Myo6), and membrane-actin linkers (Myo1b/c).

After CK-666 treatment, 23 actin-related proteins were identified (out of 166), with several overlapping hits compared to the non-treated case (Fig. 6b and Supplementary Fig. 8b). Direct comparison of all the proteins in the two conditions showed changes in Eps8’s interactome. Arpc5 disappeared upon treatment, while other Arp2/3 complex proteins had a fold change decrease tendency (Fig. 6c). Furthermore, we saw a clear change in proteins that are involved in actin assembly/disassembly. Avil and Ssh1 were lost (only found in the non-treated sample), while a greater enrichment of actin (Actb) and the Cdc42-effector proteins Cdc42ep1 (MSE55) and Cdc42bpa (MRCKα), which promote long extensions in fibroblasts (*61*) and deactivate cofilin through LIM kinase activation (*62*), respectively, were present in the CK-666-treated sample (Fig. 6c). Gene ontological (GO) analysis (Supplementary Fig. 8d, e) showed relative decreases in GO biological processes associated with the Arp2/3 complex, actin filament severing and capping, the negative regulation of actin filament polymerization, and vesicle-mediated transport; no relative changes were observed for pointed-end capping and F-actin bundle assembly. Together these data reveal a novel ability for Eps8 to switch its association with different proteins when branched actin networks are inhibited.

### IRSp53 acts as a converging platform for actin-based processes

Our data show that IRSp53 cooperates with Eps8 to facilitate processes that lead to protrusions obtaining lengths characteristic of TNTs. How these proteins coordinate with other F-actin regulators (e.g., polymerases) in this process is unclear. Therefore, we analysed IRSp53’s interactome and identified 34 and 50 actin-related protein hits in non-treated and CK-666-treated conditions, respectively (Supplementary Fig. 9a, b). From these hits, direct comparison of non-treated versus treated samples showed no significant difference (Fig. 7a) and a STRING-generated network (Fig. 7b) shows IRSp53 has an enlarged interactome as compared to Eps8. Like with Eps8, we found IRSp53 interfaces with the Arp2/3 complex and associates with proteins such as Shank3, Drebrin (Dbn1) and Lamellipodin (Raph1) that sculpt branched networks in dendritic spines and lamellipodia (*63, 64*). However, far more WAVE and WASP proteins were observed for IRSp53 (11 proteins) as compared to Eps8 (Wasf1 only). Unlike Eps8’s interactome, F-actin polymerases like the formin delphilin (Grid2ip), and those known to bind IRSp53 via SH3-mediated interactions (*28, 34*), like Vasp and Enah, were identified. IRSp53’s greater link to polymerization processes appears consistent with the observation that more actin isoforms (Actb, Actc1, Actg1, Actl6a) as well as profilin (Pfn1) were present as compared to Eps8 (Fig. 6). Furthermore, bundling proteins such as espin (Espn) (*65*) and fascin (Fscn1) were noted. Overall, these data show IRSp53 establishes a stable, multivalent interaction profile necessary to converge and promote processes affecting F-actin polymerization; however, we did not observe major changes following CK-666 treatment suggesting that Eps8’s interaction with IRSp53 (which is increased upon Arp2/3 inhibition) switches the system towards longer actin filament formation necessary for TNTs.

## Discussion

TNTs were identified as long, actin-supported protrusions mediating direct cargo transfer between distant cells (*7, 8*). This transfer function in different physiological and pathological contexts (*5, 9, 12*–*15, 18*–*22, 24, 25*) is a key difference between TNTs and filopodia (*5*). While actin is ubiquitously found in TNTs, how cells develop protrusions spanning greater distances than filopodia has remained unclear, and is a remarkable feat considering any TNT must rely on the polymerization of cytoskeletal elements far from the cell soma where globular actin (G-actin) is at a higher effective concentration. In this work we sought to address how actin-based structures, such as TNTs, are able to achieve the lengths necessary for connecting remote cells for communication and whether this process relies on unique cellular machinery.

We showed that TNT formation occurs at distances significantly higher than the average length of filopodia (Fig. 1), and that Arp2/3 inhibition favours both TNT formation (Fig. 2c) and actin polymerization over longer distances in nanotubes mechanically pulled from cells (Fig. 3). This indicates that TNTs result from a balance between linear and branched actin networks (Fig. 2 and Supplementary Fig. 4). These results bear similarity to reports in yeast, and in fibroblasts and macrophages, where the inhibition of Arp2/3, or the knockout of Arpc2, promoted formin-driven cable assembly (*66*), stress fibres and filopodia (*67, 68*), suggesting different actin networks are in homeostasis and compete for the same pool of polymerizable G-actin (*69*). As a filopodium’s steady-state length is limited by the establishment of a G-actin concentration gradient between the base and the barbed ends at the protrusion tip (*27, 70*), the formation of longer filaments here for TNTs could in part result from an increase in G-actin levels upon Arp2/3 inhibition.

Using Eps8 as an F-actin bundling protein shown to enhance TNTs (*46*) we identified a preferential interaction with the I-BAR protein IRSp53 in CAD cells (Fig. 4a) that promoted functional TNTs capable of vesicle transfer (Fig. 5), likely through a combined effect that promotes F-actin bundling and stabilizes the nanotube’s inherent curvatures. We found that Arp2/3 activity serves as an appreciable sink for the availability of Eps8 and IRSp53 (Fig. 6 and Fig. 7). Notably, Eps8 associates with Arpc1b and Arpc5L isoform-specific Arp2/3 complexes recently described as having higher branching activity (*71*). Eps8’s interaction with Arp2/3 is likely direct given the scarcity of WAVE/WASP proteins observed in our pull-down, and that diminished pull-down of complex proteins and loss of Arpc5–which tethers Arp2 and Arp3 to the mother filament for daughter filament branching (*72*)–was observed upon CK-666 treatment. This contrasts IRSp53’s SH3 domain-mediated (*37*) pull-down of far more WAVE/WASP proteins that were unaffected by CK-666 treatment. Such a scenario in which different pathways compete for reactants is equivalent to a mass action dependency and is a general concept controlling receptor tyrosine kinase signalling (*73*) to cytoskeletal network sizes (*74, 75*). Here, the reduced availability of Eps8 and IRSp53 when involved in branched networks diminishes their ability to drive outward extension of long, linear F-actin. Indeed, upon Arp2/3 inhibition, an Eps8-IRSp53 interaction was enhanced (Fig. 4a, b), existing Eps8- and IRSp53-positive protrusions further extended (Fig. 4f, g and Supplementary Fig. 6) and IRSp53 even re-localized to the plasma membrane driving new protrusion formation and extension (Fig. 4c, e). This also explains, for example, why IRSp53 overexpression alone partially inhibited the formation of functional TNTs in CAD cells (*46*), as a new steady-state favouring actin branching was likely promoted, an effect which is amplified if needed TNT-factors like Eps8 are limited within the cell.

Our results show a segregation between respective areas from where lamellipodia and TNTs emerge (Fig. 2 and Supplementary Fig. 3) suggesting spatial segregation of different actin networks is needed to form TNTs. While our study substantiates the role that competing networks have on the propensity for TNTs to form, regulation of protein availability and distribution to different pathways is needed. One possibility could involve 14-3-3 phospho-binding proteins (identified in IRSp53’s interactome, Fig. 7) which “clamp” IRSp53 in an inactive state (*76, 77*), thus preventing its membrane binding, partner associations (e.g., Eps8) and hence its downstream control of F-actin polymerization (*38*); however, CK-666 treatment showed no change in 14-3-3 protein pull-down in IRSp53’s interactome. More likely, is through Eps8’s novel ability to switch its protein and pathway associations when Arp2/3 activity is attenuated that results in linear actin growth at longer distances (Fig. 3) and promotes TNTs (Fig. 5). As observed in Eps8’s interactome changes (Fig. 6), regulation of F-actin severing could be achieved through: (1) diminished filament binding of barbed-end capping and actin-severing proteins (loss of Gsn, Avil and Flii); and (2) reinforcement of processes inactivating cofilin, for example, through the loss of cofilin-activating Slingshot phosphatases, and more LIM kinase activity and effectors (Cdc42ep1, Cdc42bpa) to deactivate cofilin. Proteins such as coronins (e.g., Coro2b identified in Eps8’s interactome, Fig. 6) directly dissociate Arp2/3 at branching points leading to network disassembly (*60, 78*) and would thus modify the frictional coupling of the actin bundle within the TNT to the cortex (*51*), diminishing retrograde flows and thus permit more growth. Similarly, the inhibition of myosin II-mediated contractility elongated microvilli through retrograde flow reduction (*79*). Thus, by attenuating processes that inhibit growth, the balance is shifted to processes that promote growth via Eps8- and IRSp53-mediated interactions to directly influence F-actin polymerization and bundling needed for TNTs. Our experimental work therefore reinforces *in vitro* reconstitution and theoretical work describing the control of F-actin network sizes through a polymerase to capping protein balance (*75, 80*). Finally, such a biphasic behaviour for Eps8 is analogous with reports showing dendritic versus linear network antagonism is dependent upon profilin (*81*), since WASP (*82*), formins (*83*) and Mena/VASP (*67, 81*) all contain profilin-binding sequences, and via the binding of tropomodulins to specific tropomyosin-decorated actin filaments (*58*).

Altogether, our study shows that the formation of TNTs depends upon common protein players and a balance between G-actin and accessory protein utilization in branched versus linear actin networks. When competing Arp2/3 networks are inhibited, this increases the availability of proteins like Eps8 and IRSp53 to exert their functions that positively lead to the extension and organization of F-actin while limiting pathways that lead to filament turnover and disassembly. While further studies are needed to elucidate the details of how Arp2/3 networks are spatially separated and disassembled, and how severing proteins are controlled to thus favour TNT growth, our work reinforces a general principle for actin network control for cellular protrusions where simple shifts in the balance between processes that inhibit growth versus those that promote growth dictate protrusion length scales.

## Materials and Methods

### Cell culture

Mouse neuronal Cath.a-differentiated (CAD) cells were used in this study and were a gift from Hubert Laude (Institut National de la Recherche Agronomique, Jouy-en-Josas, France). CAD cells were cultured in Gibco Opti-MEM + GlutaMAX (Invitrogen) supplemented with 10% fetal bovine serum (FBS; EuroBio) and 1% penicillin/streptomycin (100 u/mL final concentration; Thermo Fisher Scientific). Cells were cultured at 37°C in a humidified 5% CO2 atmosphere. Cells were routinely monitored for mycoplasma contamination and found to be negative.

### Drug treatments

Drugs used in this study include the Arp2/3 complex inhibitor CK-666 (*48*) and the formin agonist IMM-01 (*49*) and were purchased from Sigma-Aldrich. For drug treatments prior to immunoprecipitation experiments, we used 50–70% confluent CAD cells that were treated with 50 μM CK-666 for 1 hr. In the case of overnight drug treatments for TNT counting and co-culture experiments, cells first adhered for 4–6 hr and were then treated with 50 μM CK-666 or 1 μM IMM-01 for 16−18 hr. For optical tweezer experiments pre-attached cells were pre-treated for 1 hr with 50 μM CK-666. Control samples were treated with an equivalent amount of DMSO which did not exceed 0.001% v/v.

### Plasmids and lentiviral constructs

IRSp53-EGFP, IRTKS-EGFP, IRSp53-mCherry and IRTKS-mCherry were a kind gift from Pekka Lappalainen (University of Helsinki, Finland) and have been described elsewhere (*32*). GFP-Eps8-WT and the capping-deficient mutant GFP-Eps8-ΔCAP were a kind gift from Giorgio Scita (Istituto FIRC di Oncologia Molecolare, Italy) and have been described elsewhere (*45*). tdTomato-F-tractin and EGFP-F-tractin were a gift from Evelyne Coudrier (Institut Curie, France). Cytosolic GFP (pEGFP-N1) and mCherry (pmCherry-N1) vectors were purchased from Clontech. H2B-mCherry was a gift from Robert Benezra (Addgene plasmid #20972) and H2B-EBFP was a gift from Michael Davidson (Addgene plasmid #55243).

For co-expression studies, gene inserts from GFP-Eps8-WT, GFP-Eps8-ΔCAP and IRSp53-mCherry were PCR amplified using PrimeSTAR GXL DNA Polymerase (Takara Bio) and the amplicons were extracted from a 1% agarose gel (Monarch Gel Extraction Kit, NEB). Inserts were then cloned into a pCDH1 lentiviral vector, cut at EcoR1 and BamH1 restriction sites, using In-Fusion HD Cloning Plus (Takara Bio). Short Gly/Ser linkers were added between GFP and Eps8, and between IRSp53 and mCherry. Primers used in the cloning strategy include: EGFP-Xho1 forward (5′-GATTCTAGAGCTAGCGAATTCGCCACCATGGTGAGCA AGG-3′); EGFP-linker-Xho1 reverse (5′-CTCGAGGCTAGCGCTCCCACTCCCGCTTCC CTTGTACAGCTCGTCCATGCC-3′); Linker-mEps8 forward (5′-GTGGGAGCGCTAGCCTCGAGATGAATGGTCATATGTCTAACCGCTC-3′); mEps8 reverse (5′-ATCCTTCGC GGCCGCGGATCCTCAGTGGCTGCTCCCTTC-3′); IRSp53 forward (5′-GATTCTAGAGC TAGCGAATTCGCCACCATGTCTCTGTCTCGCTCAGAGG-3′); IRSp53-linker reverse (5′-CCGCTTCCGCTGTTTAAACGTCTTCCAGCCAGGGTGCG-3′); linker-mCherry forward (5′-CGTTTAAACAGCGGAAGCGGGAGTGGGAGCGTGAGCAAGGGCGAG GAG-3′); mCherry reverse (5′-TCCTTCGCGGCCGCGGATCCCTACTTGTACAGCTCGT CCATGC-3′).

### Transient transfection and stable lentiviral transduction

Plasmid transfections were performed using Lipofectamine 2000 (Thermo Fisher Scientific) according to the manufacturer’s protocol. DNA:Lipofectamine 2000 complexes at a ratio of 1:3 (μg:μL) were added in serum-free Opti-MEM media on cells plated in 6-well plates (70% confluency) and were incubated for 6 hr before removal. Transfection efficiency was verified 24 hr post-transfection and transfected cells were used for further experiments.

Lentiviral particles (LVs) were produced in HEK 293T cells cultured in Dulbecco’s Modified Eagle’s Medium (Thermo Fisher Scientific), supplemented with 10% FBS (EuroBio) and 1% penicillin/streptomycin (100 u/mL final concentration; Thermo Fisher Scientific) at 37°C in 5% CO2 humidified incubators. Cells were plated the day before transfection in T75 flasks (approx. 2 million) to achieve a 50–70% confluency the next day. Plasmids coding lentiviral components, pCMVR8.74 (Gag-Pol-Hiv1) and pMDG2 (VSV-G), and the plasmid of interest at a ratio of 4:1:4 (μg), respectively, were transfected using FuGENE HD Transfection reagent (Promega) according to the manufacturer’s protocol. After 48 hr, LVs were concentrated using LentiX-Concentrator (Takara Bio) and the pellet was resuspended up to 1 mL in serum-free Opti-MEM. CAD cells were plated the day before infection in 6-well plates (approx. 100,000 cells) to achieve a 50–70% confluency the next day. Cells were sequentially transduced with 100 μL of the desired LVs, each for 24 hr. After transduction, co-expressing cells were treated with 2 μg/mL puromycin for 3 days. Double-positive cells were sorted using a BD FACSAriaIII cell sorter (BD Biosciences).

### Surface micropatterning

Micropatterning on glass coverslips was performed using an adapted method based Azioune et al. (*84*). Round glass coverslips (25 mm, No. 1.5, Marienfeld) were sequentially cleaned in a bath sonicator with 70% ethanol for 10 min and 2% Hellmanex (Hellma Analytics) for 20–30 min at 30°C, and then in 1 M KOH for 20 min; copious rinsing using ultra-pure water (resistivity >18 MΩ·cm) from a Milli-Q Advantage A-10 purification system (EMD Millipore) was performed after each step. The coverslips were then dried under a stream of nitrogen and then plasma activated (Harrick Plasma) for 5 min and then functionalized overnight in a 3% v/v solution of (3-aminopropyl)trimethoxysilane (APTMS) in acidified methanol (5% v/v acetic acid). Following aminosilanization, coverslips were briefly rinsed with 70% ethanol and then annealed in a 120°C oven for 1 hr. Individual coverslips were then placed on a 60–70 μL drop of a 100 mg/mL solution of mPEG-succinimidyl valerate (MW 5000, Laysan Bio) dissolved in 0.1 M NaHCO3, pH 8.5 and left to react overnight in a humified environment to prevent evaporation. The following day, PEG-coated coverslips were rinsed in ultra-pure water and stored dry in a desiccator until they were used for deep-UV micropatterning.

Custom micropattern designs were drawn in CleWin (WieWeb) and the chrome-quartz photomasks were manufactured at Delta Mask (Enschede, NL). The PEG-coated coverslips were held in contact with the photomask using a 2.5 μL drop of water (providing an estimated spacing of 5 μm), and the photomask was placed 5 cm away (∼10 mW/cm^2^) from the low-pressure mercury lamps (Heraeus Noblelight GmbH, NIQ 60/35 XL) housed within a UV box (UVO Cleaner, Jelight); coverslips were then exposed for 5 min to deep UV light (< 200 nm) through the photomask, burning exposed regions of the PEG layer to create the desired pattern on the coverslip. Subsequently, to allow for stronger covalent binding of fibronectin to the patterned areas, *N*-hydroxysulfosuccinimide (sulfo-NHS)/1-Ethyl-3-(3-dimethylaminopropyl)carbodiimide (EDC) chemistry for amine coupling of proteins to the surface was employed. Sulfo-NHS and EDC were dissolved in 0.1 M MES, 0.5 M NaCl, pH 6.0 at a concentration of 6 and 124 μM, respectively, and the patterned sides of the coverslips were incubated with the sulfo-NHS/EDC solution for 30 min. Coverslips were then rinsed with ultra-pure water, and then incubated for 1 hr with a fibronectin (FN; Bovine plasma, Sigma-Aldrich) solution having a total concentration of 35 μg/mL (dissolved in 0.1 M NaHCO3, pH 8.5). The FN solution was a mixture of unlabeled and dye-labeled FN, where for example, fluorescent FN typically comprised 15% of the total FN concentration. The micropatterned surfaces were stored at 4°C in PBS containing 1% penicillin/streptomycin until used for cell seeding and typically used within one week.

### Fluorescent fibronectin

Rhodamine-labelled FN was purchased from Cytoskeleton. Labelling of FN with Alexa Fluor 405 succinimidyl ester (Thermo Fisher Scientific) was performed in house. Briefly, lyophilized FN (Bovine plasma, Sigma-Aldrich) was dissolved in PBS and dialyzed overnight against PBS to remove free amines. The dialyzed FN solution was then labelled at a dye to protein ratio of 70:1 (mol:mol) for 1 hr at room temperature (RT) and in the dark to protect the reaction from light. Extensive dialysis against PBS was then performed to remove free Alexa Fluor dye. The labelled FN was concentrated using Amicon Ultra centrifugal filters (Merck), and aliquoted and stored at - 80°C before use. The labelling efficiency for Alexa 405 FN was determined to be 30.

### Characterization and analysis of cell adherence on micropatterns

Reflection interference contrast microscopy (RICM) was performed on an inverted Eclipse Ti-E inverted microscope (Nikon) equipped with an Antiflex 63X 1.25 NA oil immersion objective (Zeiss Plan-Neofluar) and a Moment sCMOS camera (Teledyne Photometrics). Light from a mercury-vapor arc lamp was bandpass filtered (546.1/10 nm, Melles Griot) and then directed towards a filter cube containing a 50/50 beam splitter and two crossed polarizers prior to entering the objective. An aliquot of suspended CAD cells (∼20,000 cells) was added on top of a micropatterned substrate mounted on a Chemlide CMB 35 mm magnetic chamber (Quorum Technologies). Acquisition was controlled through MetaMorph and brightfield and RICM images were acquired every 60 sec over 5 hr to capture as cells begin to adhere and spread on the micropatterns. A reference fluorescence image of the FN micropatterns was taken at the first timepoint only. RICM images were variance filtered in ImageJ/Fiji to highlight the contour of the cell as the surrounding background exhibited low signal variance. Ten variance-filtered images were then used as a training dataset in Weka for automatic segmentation of the contour of adhered cells. Segmented images were binarized and the area was determined in ImageJ/Fiji through time.

### Tracking cell confinement on micropatterns

Brightfield and fluorescence imaging was performed on an inverted Eclipse Ti-E or Ti2 inverted microscope (Nikon) equipped with either a CSU-X1 or a CSU-W1 spinning disk confocal scanning unit (Yokogawa), a fibre-coupled Coherent OBIS laser illumination system (405/488/561/640 nm), a 40X 1.15 NA objective (CFI Apochromat LWD Lambda S, Nikon), and a Prime 95B sCMOS camera (Teledyne Photometrics). CAD cells were first pre-adhered for 1 hr to the micropatterns and then imaged every 5 min over 18 hr using MetaMorph or NIS-Elements (Nikon) for instrumental control. Reference fluorescence images of the FN micropatterns were taken at the first timepoint only. Cells in the brightfield images were segmented from the background using an Ilastik-trained model resultant from a four-image training dataset. Resulting binarized images were then inputted into TrackMate to track an individual cell’s centre-of-mass, with respect to the centre of the cell’s micropattern, through the acquisition. Tracks were overlaid on images using “Track Manager” in Icy.

### Bead preparation for nanotube pulling experiments

Streptavidin-coated polystyrene beads (3 μm in diameter, SVP-30-5, 0.5% w/v, Spherotech) were washed three times in a 10X volume of PBS and recovered by centrifugation (12,000 rpm, 5 min). Beads were then resuspended in PBS to a concentration of 0.05% w/v, and an appropriate amount of a 1 mg mL^−1^ biotin-conjugated concanavalin A (ConA) solution (C2272, Sigma-Aldrich) was added to the bead suspension assuming a binding capacity of 10 μg ConA per mg of beads. The mixture was incubated overnight at 4°C on a tabletop shaker. ConA-coated beads were rinsed three times according to the steps above and finally resuspended in PBS to a concentration of 0.5% w/v. ConA-beads were stored at 4°C and generally usable up to one month.

### Nanotube pulling experiments

A home-built optical tweezer (OT) coupled to an inverted Eclipse Ti C1 Plus confocal microscope (Nikon) was used as previously described (*38*). Briefly, a 3-watt 1064 nm Ytterbium fiber laser (IPG Photonics) was expanded through a Keplerian telescope and directed towards the back aperture of a 100X 1.45 NA oil immersion objective (Nikon CFI Plan Apochromat Lambda). The viscous drag method, including Faxen’s correction for calibration near surfaces, was used to determine the OT trap stiffness which averaged 60 pN μm^−1^. Displacements of the bead from the fixed trap centre were recorded on a CCD camera (Marlin F-046B, Allied-Vision) at a frame rate of 20 fps and videos were later analysed using a custom MATLAB (Mathworks) script based on the “imfindcircles” function to determine the bead’s centre of mass. Forces were calculated from the bead positions according to the equation *F* = *κ* · Δ*x* where *κ* is the trap stiffness and Δ*x* is the displacement of the bead from its initial reference position determined before nanotube pulling. A Nano-LP100 piezo-driven stage (MadCityLabs) was used for lateral XY movements on the setup. A temperature and CO2 controllable stage-top incubator (STXG-WELSX, Tokai Hit) maintained cells at 37°C in a humidified, 5% CO2 atmosphere during experimentation.

The day before experimentation, CAD cells expressing EGFP-F-Tractin were seeded on micropatterned coverslips in which the fibronectin patches were separated at a distance of 50 μm to ensure cells were well-separated for micromanipulation. One-hour prior to experimentation the phenol-containing culture medium was removed, cells were rinsed with PBS, and exchanged for a phenol-free Opti-MEM medium containing: 10% FCS, ProLong™ Live Antifade Reagent (Invitrogen) at a 1:75 dilution, 2 mg mL^−1^ β-Casein (>98% pure, from bovine milk, Sigma-Aldrich) for surface passivation, and 50 μM CK-666 (or the equivalent volume of DMSO for control experiments). The cells were taken to the optical tweezer setup and labeled with Cell Mask™ Deep Red plasma membrane stain (Invitrogen) at a 1:2000 dilution for 10 min, and ConA-coated beads were added (1:50–1:100 dilution). Using a custom LabVIEW (National Instruments) program to control the piezo stage, membrane nanotubes were pulled by trapping an isolated floating bead, bringing it into contact with the cell for a short period of time (<10 sec), and then moving the cell away from the bead in the X direction. Confocal image acquisition using 488 nm (Coherent) and 642 nm (Melles Griot) lasers were controlled with the EZ-C1 software (Nikon). Fluorescence was bandpass filtered (ET525/50, ET665, Chroma) and detected using τ-SPAD single-photon avalanche diodes (PicoQuant) that were controlled by the SymPhoTime 64 software (PicoQuant). Images encompassing the nanotube and some of the cell body (typically 1024 × 512 pixels, 5X zoom) were gathered every 30 sec after pulling.

### Quantification of actin profiles in pulled nanotubes

Image analysis was performed using custom written macros in ImageJ/Fiji. Images for the membrane and actin channels were first background subtracted, and the actin channel was then normalized by the average cytosolic signal measured within the cell interior to account for differences in F-Tractin expression. From the membrane channel image, the nanotube’s cross-section was fit to a Gaussian function to determine the ±2σ width of the Gaussian profile. Using the line selection tool, a line was drawn starting within the cytosol, intersecting through the plasma membrane and then following all along the length of the nanotube; the width of the line was set to the measured ±2σ width in order to encompass the entire thickness of the nanotube. Profiles were then extracted and the actin intensity plotted as a function of the length of the nanotube *X*, where *X* = 0 was defined as the position of the max intensity value found within the plasma membrane rim near the base of the nanotube. For curve averaging, profiles were interpolated in MATLAB to obtain vectors of all the same length.

### GFP-trap immunoprecipitation

GFP-trap was performed using anti-GFP nanobody-coated agarose beads (Chromotek) and was carried out following the supplier’s recommendations. Briefly, 1.2 million CAD cells expressing the GFP-tagged “bait” protein of interest were plated in 100 mm dishes. The subsequent day the cells were briefly washed with PBS after drug treatment, and then scraped and centrifuged at 500 g for 3 min at 4°C. The supernatant was gently removed, and lysis buffer was added (4–5 volumes of lysis buffer for 1 volume of cell pellet). The lysis buffer consisted of 50 mM Tris, 1% Triton, 300 mM NaCl, 5 mM MgCl2, cOmplete™ Mini EDTA-free Protease Inhibitor Cocktail (Merck) at pH 7.4. The cells were lysed on ice for 30 min, after which the lysate was cleared at 14,000 rpm for 30 min at 4°C. The cell lysate protein concentration was quantified using a Bradford protein assay (Bio-Rad). Agarose beads were rinsed and equilibrated in PBS, recovered by centrifugation and then resuspended in PBS +/+ (1:1 ratio). After which 20 μL of resuspended beads were added per immunoprecipitation (IP) sample (300 μg of total protein) along with 300 μL of dilution buffer (10 mM Tris, 150 mM NaCl, 0.5% Triton, pH 7.5). The IP mixture was diluted with sterile distilled water, so that the final Triton concentration did not exceed 0.5% v/v, and was then incubated for 1 hr at 4°C with gentle agitation to bind proteins. Three washes of 3 min each were carried out with wash buffer (10 mM Tris, 150 mM NaCl, 0.5% Triton, pH 7.5). Finally, bound proteins were eluted with 20 μL of 2X Laemmli sample buffer (Bio-Rad) and the sample was subsequently denatured at 100°C for 5 min.

### SDS-PAGE and Western Blot analysis

SDS-PAGE was performed in 1X MOPS buffer (Sigma-Aldrich) on Criterion 4–12% precast gels (Bio-Rad). Wells were loaded with the total eluent volume from the IP samples, containing 300 μg of protein, whereas 10% (30 μg) was loaded for the lysate control samples (input). The migration was done at 120 mV for approximately 2 hr. Proteins were transferred (100 V, 1 hr) onto PVDF (polyvinylidene difluoride) membranes (GE Healthcare Life Sciences) using 1X Tris/Glycine buffer (in-house 50X solution containing 250 mM Tris and 1920 mM Glycine). After transfer, the membrane was blocked using 5% non-fat milk in 0.1% TBS-T (1X Tris-Buffered Saline, 0.1% Tween 20, Bio-Rad) for 1 hr at RT. Primary antibodies were added at the appropriate dilution, and membranes were incubated overnight at 4°C under agitation. Primary antibodies used include: rabbit polyclonal anti-GFP (1:4000, A-6455, Thermo Fisher Scientific), rabbit polyclonal anti-IRSp53 (1:1000, HPA023310, Sigma-Aldrich), rabbit polyclonal anti-IRTKS (1:2000, HPA019484, Sigma-Aldrich), and mouse monoclonal anti-α tubulin (1:5000, T5168, Sigma-Aldrich) as a loading control. Membranes were washed three times in TBS-T for 10 min and mouse and rabbit horseradish peroxidase (HRP)-tagged secondary antibodies were added to the membrane at a 1:5000 dilution in 5% milk, for 1 hr at RT. Membranes were revealed using Amersham ECL Prime (Cytiva) chemiluminescent detection reagent and imaged using an ImageQuant LAS 500TM imager (GE Healthcare Life Sciences).

To quantify relative changes in band intensities (measured as an average value), regions of interest of the same size were drawn in ImageJ/Fiji to encompass the appropriate GFP band of the bait protein (indicates the amount of eluted bait protein), the band of the prey protein, and background values adjacent to the measured bands. Background-corrected band intensities were finally normalized by taking the prey band divided by the bait band.

### TNT quantification

#### On micropatterns

CAD cells were mechanically detached in serum-free Opti-MEM containing 0.5 mM EGTA (to prevent cadherin-mediated cell clumping), strained through a 40 μm nylon cell strainer (Corning) and diluted to a concentration of 175,000 cells/mL using Opti-MEM containing 10% FBS to better ensure singularized cells. Micropatterned coverslips were generally attached to the bottom of open 35 mm dishes (P35G-1.5-20-C, MatTek) using silicon grease (Dow Corning), and 60,000 cells were seeded within the volume defined by the 20 mm dish opening. Otherwise, micropatterned coverslips were placed in 6-well plates and 250,000 cells were used for seeding in this case. After 4–6 hr of adherence, unattached cells were then removed manually using a pipette by gently rinsing with PBS. Cells were then cultured 16−18 hr overnight prior to a two-step fixation and dye staining.

Cells were fixed with 2% paraformaldehyde (16% PFA aqueous solution, methanol free, EM grade, Electron Micorscopy Sciences), 0.05% glutaraldehyde (25% GA aqueous solution, Grade I, Sigma-Aldrich), 0.2 M HEPES (1 M HEPES, Gibco) in PBS for 20 min at 37°C, followed by a second fixation in 4% PFA, 0.2 M HEPES in PBS for 20 min at 37°C. Following fixation, cells were quenched in 50 mM NH4Cl for 10–30 min at RT and carefully washed in PBS prior to dye labelling. Alexa Fluor 647 phalloidin or Rhodamine phalloidin (1:50–1:200 dilution in PBS, 30 min incubation in the dark at RT; Thermo Fisher Scientific) was used for F-actin staining, WGA Alexa Fluor 488 conjugate (1:300 dilution in PBS, 20 min incubation in the dark at RT; Thermo Fisher Scientific) was used to stain the plasma membrane, and 4’,6-diamidino-2-phenylindole (DAPI) (1:1000 dilution, 2 min incubation in the dark at RT) was used to stain the nuclei. Between staining, samples were washed with PBS. Finally, samples were mounted/sealed using Aqua-Poly/Mount (Polysciences) or Mowiol 4-88 (Sigma-Aldrich) and left to polymerize overnight in the dark.

Samples were imaged using a 40X 1.3 NA oil immersion objective (Zeiss Plan Apo) on an inverted confocal microscope (Zeiss LSM 700) equipped with 405/488/561/640 nm lasers and controlled by the ZEN software. The whole cellular volume, typically 12 μm in thickness, was imaged by acquiring 0.4 μm-thick slices. Images were analyzed in the Icy software using the “Manual TNT annotation” plug-in (http://icy.bioimageanalysis.org/plugin/manual-tnt-annotation/) for manual identification and numeration of cells connected by TNTs. Connections between cells on neighbouring micropatterns were annotated as TNTs if the following criteria were met: (i) they were thin (diameter < 800 nm), (ii) membranous and F-actin positive, (iii) did not contact the substrate and thus would be found in the middle and upper stacks, and (iv) were continuous along their length without obvious breaks (e.g., tip-to-tip bound filopodia) to directly link the cell bodies of the connected cells. The percent of TNT-connected cells on micropatterns was obtained by counting the TNT-connected cells and dividing by the number of nearest-neighbour cell-occupied micropatterns (i.e., isolated cell-occupied micropatterns were excluded since nearest-neighbour micropatterns are unoccupied and thus incapable of having a TNT spanning between micropatterns).

#### In normal cell culture

In Supplementary Figs 4 and 5, 220,000 cells were plated into 35 mm Ibidi μ-dishes (Ibidi), cultured 18 hr overnight and then fixed, stained and imaged similarly to what was described above for the micropatterned samples. The percent of TNT-connected cells was obtained by counting the TNT-connected cells in an image and dividing by the total number of cells present in the field of view.

### Manual single-cell analysis of TNT origin

Wild-type CAD cells plated on D*15* micropatterns (see TNT quantification on micropatterns, Fig. 2) were classified into either a mixed cell phenotype (exhibiting hairy/filopodial and lamellipodial/ruffled characteristics) or a lamellipodia-only phenotype. Then, the immediate cellular vicinity from which the TNT was originating was categorised. TNTs were categorised as emanating from a filopodia-rich region, directly from a lamellipodia/ruffle region, or spatially separated above a lamellipodia/ruffle region.

### Quantification of vesicle transfer by co-culture assays

#### By confocal microscopy

Vesicle transfer assays such as those in Supplementary Fig. 2 and Fig. 5 were performed on *D*15 micropatterns, while IMM-01 drug-treated co-cultures were performed in 35 mm Ibidi μ-dishes (Supplementary Fig. 4). Donor cells were stained with DiD (Thermo Fisher Scientific; 1:3000 dilution in culture media for 30 min in the dark at 37°C) to visualize vesicles, whereas cells expressing a fluorescently tagged histone 2b (H2B) fusion protein were used as acceptor cells. Predominately, H2B-EBFP-transfected CAD cells were used as an acceptor cell population unless otherwise stated. Donors and acceptors were co-cultured overnight (16−18 hr) at a 1:1 ratio; the total amount of cells plated in Ibidi μ-dishes was 200,000, while the number of cells plated on *D*15 micropatterns (placed in 6-well plates) was 250,000. In parallel, supernatant controls were performed in Ibidi μ-dishes to measure secretion-based vesicle transfer. Here, equal amounts of donor and acceptor cells (100,000 for Ibidi-based co-cultures and 125,000 for micropattern-based co-cultures) were separately cultured. After 16−18 hr, the conditioned media of the donors was collected, briefly centrifuged (1,000 rpm, 4 min) to remove dead cells, and then exchanged for the media of the acceptor cell sample. The acceptors cells were then cultured for another overnight incubation with the conditioned media. All samples were fixed, stained and imaged according to the protocol described above for TNT quantification. The Icy software was used to annotate acceptor cells with and without internalized DiD-positive vesicles. The percent of total transfer was assessed by quantifying the number of acceptor cells containing donor-derived DiD-vesicles and dividing it by the total number of acceptor cells analysed across the acquired images for a given sample.

#### By flow cytometry

In Supplementary Fig. 5d, CAD cells expressing IRTKS-EGFP where stained with DiD (donor population) and cultured at a 1:1 ratio (300,000 total cells) with H2B-mCherry expressing CAD cells (acceptor population) in 6-well plates. In parallel, co-cultures using EGFP expressing donor cells were used as controls. After an overnight co-culture (16−18 hr), cells were mechanically detached and washed briefly with a dilute trypsin solution (0.03% w/v trypsin in PBS) to remove non-internalized vesicles attached on the cell exterior. Cells were passed through a 40 μm nylon cell strainer to separate cell aggregates and fixed in 2% PFA. Flow cytometry data was acquired with a FACSymphony A5 flow cytometer (BD Biosciences). GFP, mCherry and DiD fluorescence were analysed at excitation wavelengths of 488, 561 and 647 nm, respectively. Ten thousand events were acquired for each condition and data was analysed using FlowJo (BD Biosciences). The percent of total transfer was calculated by dividing the number of double positive cells (H2B-mCherry acceptor cells containing donor-derived DiD-vesicles) with the total number of analysed H2B-mCherry acceptor cells (Q2/Q1). Two biological repeats were conducted with each condition done in triplicate.

### Time-lapse microscopy of protrusion formation

CAD cells at 50–70% confluency were co-transfected with IRSp53-EGFP and tdTomato-F-tractin or GFP-Eps8-ΔCAP and tdTomato-F-tractin according to the transfection protocol above. Six hours prior to the acquisition, 220,000 of the co-transfected cells were plated in 35 mm dishes (Ibidi), or 250,000 cells were plated on 40 μm fibronectin micropatterns placed in 6-well plates (Supplementary Fig. 6, Example 3). Samples were mounted on a Chemlide CMB 35 mm magnetic chamber (Quorum Technologies). Time-lapse imaging was performed on an Eclipse Ti-E inverted microscope (Nikon) equipped with a CSU-X1 spinning disk confocal scanning unit (Yokogawa), a 60X 1.4 NA oil immersion objective (Nikon Plan Apo VC), a fiber-coupled Coherent OBIS laser illumination system (405/488/561/640 nm), and Prime 95B sCMOS cameras (Teledyne Photometrics) for single or dual camera mode acquisition. Acqusition was controlled by MetaMorph. Two positions with z-stack slices (0.6 μm thickness) covering the 12 μm cell volume were acquired at a 1-min interval over 90 min (except when imaging IRSp53 recruitment at the membrane which was imaged with a 30-sec interval). Cells were imaged 30 min prior to flowing in CK-666 at a final concentration of 50 μM, that was subsequently imaged for 60 min. A humidified 37°C and a 5% CO2 environment was maintained in a Life Imaging Services Gmbh enclosure.

### Filopodia quantification and length measurements

For fixed conditions (Fig. 1g), filopodia were quantified using the automated analysis pipeline developed by Bagonis et al. (*85*). Maximum intensity projection images of filopodia (both surface- and non-attached) were created and the cell bodies were manually segmented for filopodia detection using phalloidin staining. When needed, Hyugens (Scientific Volume Imaging) was used for image deconvolution to improve filopodia detection in samples having weak actin fluorescence.

For time-lapse imaging (Fig. 4d, g), maximum intensity projection images were created in ImageJ/Fiji. In the first 30 min of imaging (control condition prior to CK-666 addition), a protrusion containing actin (as F-tractin) and the protein of interest was followed and when it reached its maximum length (e.g., just before retraction), its length was manually measured from its base to the tip. The same procedure was performed after CK-666 treatment, with the difference being that the imaging duration was increased to 60 min in order to sample length changes in manually tracked protrusions.

### Fluorescence profile measurements

The intensity of IRSp53 at the membrane (Fig. 4e) before and after the addition of CK-666 was analysed in ImageJ/Fiji (“Plot Profile” plugin) by drawing a straight line (10 μm long) perpendicular to the cell edge. The middle of the line was positioned directly at the boundary of cell edge and the background such that individual curves could be properly aligned and overlaid.

### Proteomic analysis by mass spectrometry

#### Sample preparation

CAD cells were plated in 100 mm dishes (800,000 cells) the day before transfection. Following a 24-hr transfection, GFP-Eps8-WT or IRSp53-EGFP cells, along with cytosolic GFP-expressing cells, were treated in parallel either with DMSO or with 50 μM CK-666 for 1 hr. After washing and lysing the cells, samples were processed for GFP-trap immunoprecipitation (described above). After protein binding, two 3-min washes were done in lysis buffer followed by three 3-min washes in the dilution buffer prior to on-bead digestion and mass spectrometry (MS) analysis. Three independent preparations were performed for Eps8, while five independent preparations were performed for IRSp53.

#### On-bead digestion followed by LC-MS/MS analysis

Digestion was performed strictly as described by Chromotek. Briefly, beads were resuspended in digestion buffer (50 mM Tris-HCl pH 7.5, 2 M urea, 1 mM DTT and 5 μg/μl of trypsin (Promega)) for 3 min at 30°C. Supernatants were transferred to new vials and beads were washed twice using 50 mM Tris-HCl pH 7.5, 2 M urea and 5 mM iodoacetamide buffer. All washes were pooled and incubated at 32°C for overnight digestion in the dark. Peptides were purified using a standard C18-based clean-up protocol using a Bravo AssayMap device (Agilent). LC-MS/MS analysis of digested peptides was performed on an Orbitrap Q Exactive Plus mass spectrometer (Thermo Fisher Scientific) coupled to an EASY-nLC 1200 (Thermo Fisher Scientific). A home-made C18 column was used for peptide separation and consisted of a 30 cm nano-HPLC capillary column (75 μm inner diameter) filled with 1.9 μm Reprosil-Pur Basic C18-HD resin (Dr. Maisch HPLC GmbH) that terminated with a silica PicoTip^®^ emitter tip (New Objective). The column was equilibrated and peptides were loaded in 100% solvent A (H2O, 0.1% formic acid (FA)) at 900 bars. Peptides were eluted at 300 nL min^−1^ using a linear gradient of solvent B (acetonitrile (ACN), 0.1 % FA) from 2% to 35% during 55 min, 35% to 60% during 10 min, and 60% to 90% during 5 min (total chromatographic run was 80 min including a high ACN step and a column regeneration step). Mass spectra were acquired in the data-dependent acquisition mode with the XCalibur 2.2 software (Thermo Fisher Scientific). Automatic switching between MS and MS/MS scans was performed to select the 10 most abundant precursor ions (top10 method) from the survey scan for higher-energy collision dissociation (HCD) peptide fragmentation. Survey MS scans were acquired at a resolution of 70,000 (at m/z 400) with a target value of 3 × 10^6^ ions and was limited in range from 400 to 1700 m/z. HCD peptide fragmentation was performed with a normalized collision energy (NCE) set at 27%. Intensity threshold for ion selection was set at 1 × 10^6^ ions with a charge-state exclusion of z = 1 and z > 7. The MS/MS spectra were acquired at a resolution of 17,500 (at m/z 400). Isolation window was set at 1.6 Th. Dynamic exclusion was employed within 30 s.

#### Protein identification and quantification

MS Data were searched using MaxQuant (version 1.6.6.0) employing the Andromeda search engine (*86*) against a reference *Mus musculus* proteome (55,470 entries; downloaded from Uniprot June 1, 2021). The following search parameters were applied: carbamidomethylation of cysteines was set as a fixed modification, while oxidation of methionine and protein N-terminal acetylation were set as variable modifications. The mass tolerances in MS and MS/MS were set to 5 ppm and 20 ppm, respectively. Maximum peptide charge was set to 7 and 5 amino acids were required as a minimum peptide length. At least 2 peptides (including 1 unique peptide) were asked to report a protein identification. A false discovery rate (FDR) of 1% was set for both protein and peptide levels. Intensity-based absolute quantification (iBAQ) values were generated from the sum of peak intensities of all peptides corresponding to a specific protein divided by the number of observable peptides. The match between runs features was allowed for biological replicate only.

#### Statistical analysis

Proteins identified in the reverse and contaminant databases and proteins “only identified by site” were first removed from the analysis. Remaining proteins with at least 2 peptides were kept for further analysis and the associated peptide intensities were summed and log transformed (log2). Summed intensity values were normalized by median centering within conditions (*normalizeD* function of the R package *DAPAR*) (*87*). Proteins without any iBAQ value in a condition, while present in another, have been considered as proteins quantitatively present in a condition and absent in the other. Next, missing values were imputed using the *impute*.*mle* function of the R package *imp4p* (*88*). Proteins with a fold-change inferior to 1.5 (i.e., log2(FC) < 0.58) are considered not significantly differentially abundant between conditions. Statistical testing of proteins with a FC superior to 1.5 was conducted using a limma t-test thanks to the R package *limma* (*89*). An adaptive Benjamini-Hochberg procedure was applied on the resulting p-values thanks to the *adjust*.*p* function of the R package *cp4p* (*90*) using the robust method described in (*91*) to estimate the proportion of true null hypotheses among the set of statistical tests. The proteins associated to an adjusted P-value inferior to an FDR level of 1% have been considered as significantly differentially abundant proteins. Thus, proteins of interest are therefore those considered to be present in one condition and absent in another, and those proteins identified based on pairwise comparison of intensities according to the adjusted P-value and FC thresholds.

### Bioinformatic analysis

UniProt identifiers/accession numbers of the proteins of interest (specifically those related to the actin cytoskeleton) were inputted into the online Search Tool for Recurring Instances of Neighbouring Genes (STRING) database (*54*) to generate an interaction network visualized using Cytoscape. Medium confidence (minimum score, 0.4) interactions were sourced from databases, experiments, co-expression, neighborhood and gene fusion evidence. The proteins in the network were grouped manually according to their function and the networks were then exported as .svg files for color coding in Adobe Illustrator for easier visualization. Gene Ontology (GO) analysis of biological processes was performed using PANTHER 17.0, employing Fisher’s Exact test with a false discovery rate (FDR) correction. The whole *Mus musculus* gene database was used as a reference list. Output parameters for a given GO term include: 1) the protein count in the input; 2) a reference count (Ref cout) for the total number of proteins in the mouse database assigned to a given GO term; 3) the fold enrichment value (observed/expected) which is determined from the protein count divided by the number of proteins expected to be annotated in a randomly generated network of the same size; and 4) the FDR that measures the significance of the fold enrichment.

### Statistical analysis

Statistical analysis was performed in GraphPad Prism. P < 0.05 was considered statistically significant and all P values are indicated in the figures. The specific statistical tests performed, the number of independent experiments and the total number of samples analysed are indicated in the figure captions.

## Data availability

The mass spectrometry proteomics data have been deposited to the ProteomeXchange Consortium via the PRIDE (*92*) partner repository with the dataset identifier PXD035976. All data needed to evaluate the conclusions in the paper are present in the paper and/or the Supplementary Materials.

## Acknowledgements

We thank all laboratory members for insightful discussions, particularly Christel Brou, Stéphanie Lebrenton and Feng-Ching Tsai for their critical reading of the manuscript. We are also grateful for the administrative support provided by Reine Bouyssie (Membrane Traffic and Pathogenesis Unit, Institut Pasteur). We additionally thank Juliette Wu, a former intern, for help in some early immunoprecipitation experiments, and Fanny Tabarin-Cayrac (Institut Curie) and Pierre-Henri Commere (Institut Pasteur) for cell sorting assistance. The authors greatly acknowledge experimental support from the Institut Pasteur Unit of Technology and Service for Cytometry and Biomarkers (UTechS CB), and the Photonic BioImaging platform (UTechS PBI) which is a member of the France BioImaging infrastructure (FBI) network that is supported by the French Agence Nationale de la Recherche (ANR-10-INBS-04). J.M.H. was supported by a Pasteur Foundation Postdoctoral Fellowship and N.L. was supported by the Sorbonne Université (doctoral grant number 3210/2018). P.B. acknowledges support from the Centre National de la Recherche Scientifique (CNRS Défi Instrumentation aux limites 2018), and is a member of the CNRS consortium Approches Quantitatives du Vivant (AQV), Labex Cell(n)Scale (ANR-11-LABX0038) and Paris Sciences et Lettres (ANR-10-IDEX-0001-02). C.Z. acknowledges support from the Equipe Fondation Recherche Médicale (FRM EQU202103012692). This work was supported by grants from the Université Paris Sciences et Lettres-QLife Institute (ANR-17-CONV-0005 Q-LIFE) and the Agence Nationale de la Recherche (ANR-20-CE13-0032 LiveTuneL) to P.B. and C.Z.

## Contributions

J.M.H., P.B. and C.Z. conceived the study and acquired funding. J.M.H. and N.L. designed and performed experiments, analysed the resulting data, and wrote the original manuscript. T.C. performed mass spectrometry measurements and performed statistical analyses with Q.G.G under the supervision of M.M. D.C. performed IMM-01 experiments. A.B. provided molecular biology expertise, reagents and generated lentiviral constructs used in the study. S.D. provided technical expertise and design assistance for the micropatterning protocol. J.M.H., N.L., P.B. and C.Z. reviewed and edited the final manuscript. All authors approved the manuscript prior to submission.

