## Supplementary Information for "A balance between actin and Eps8/IRSp53 utilization in branched versus linear actin networks determines tunneling nanotube formation"

---

---

**This pdf includes:**

Supplementary Figs. 1 to 9
Supplementary Videos 1 to 12

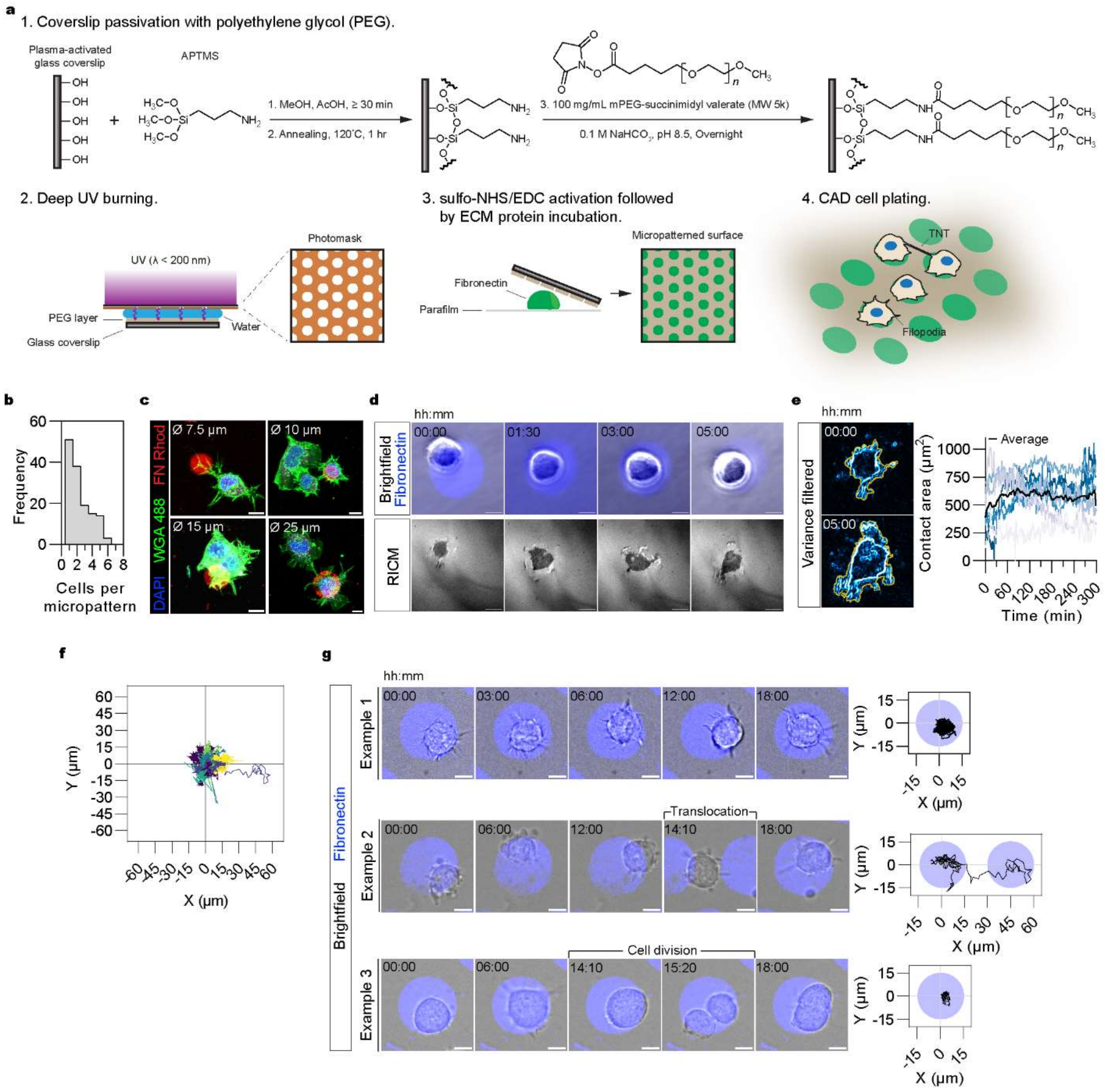


**Supplementary Fig. 1: Micropatterning approach for assessing TNT formation.**
(**a**) Scheme depicting the fabrication process of the micropatterned surfaces, including (1) coverslip passivation with covalently attached PEG molecules, (2) deep UV printing using a chrome-quartz photomask, (3) covalent attachment of fibronectin (FN) using N-sulfohydroxysuccinimide (sulfo-NHS)/1-Ethyl-3-(3-dimethylaminopropyl)carbodiimide (EDC) amine coupling chemistry, and finally (4) CAD cell plating to better discriminate TNT-connected cells from morphologically similar and shorter filopodia. (**b, c**) Micropatterns having a diameter (Ø) of 31 µm ($A$ ~ 750 µm$^2$), which permitted more than one cell to adhere per micropattern on average, were the most optimal for CAD cell adherence and their long-term immobilization, as decreasing diameters resulted in poorer cell patterning and frequent cell infiltration in the surrounding PEG region. (**b**) Histogram of the number of cells per micropattern following overnight culture ($n$ = 140 micropatterns). (**c**) Screen of different micropattern diameters for CAD

cell adherence. CAD cells exhibited poorer micropatterning at Ø < 31 μm (blue, DNA labelling with DAPI; green, membrane labelling with wheat germ agglutinin (WGA); red, FN Rhodamine). (**d, e**) Reflection interference contrast microscopy (RICM) was used to measure the contact area of CAD cells through time as a proxy for their surface adhesion. (**d**) Representative brightfield and RICM images of early CAD cell adherence (over 5 hours) on FN micropatterns 31 μm in diameter (blue, FN Rhodamine). The darker contrast in RICM images indicates shorter distances between the glass surface and the adhering plasma membrane of the cell. (**e**) Left: RICM images were subjected to a variance filter and then segmented using a Weka-trained model to obtain the adhered contact area of the cell (yellow contours). Right: Plot of the contact area measured over time ($n$ = 7 cells). The average contact area begins to plateau near 60 minutes (and remains stable afterwards) indicating cells have formed stable adhesions to the FN micropatterns. (**f, g**) Assessment of CAD cell confinement over an extended period of time (18 hrs). (**f**) Plot of individual cell trajectories normalized with respect to the centre of their micropattern (origin) ($n$ = 56 cell trajectories). (**g**) Representative time-lapse images of CAD cells and their corresponding trajectories. Fibronectin micropatterns are false-coloured in blue (rhodamine-labelled fibronectin). Scale bars, 10 μm.

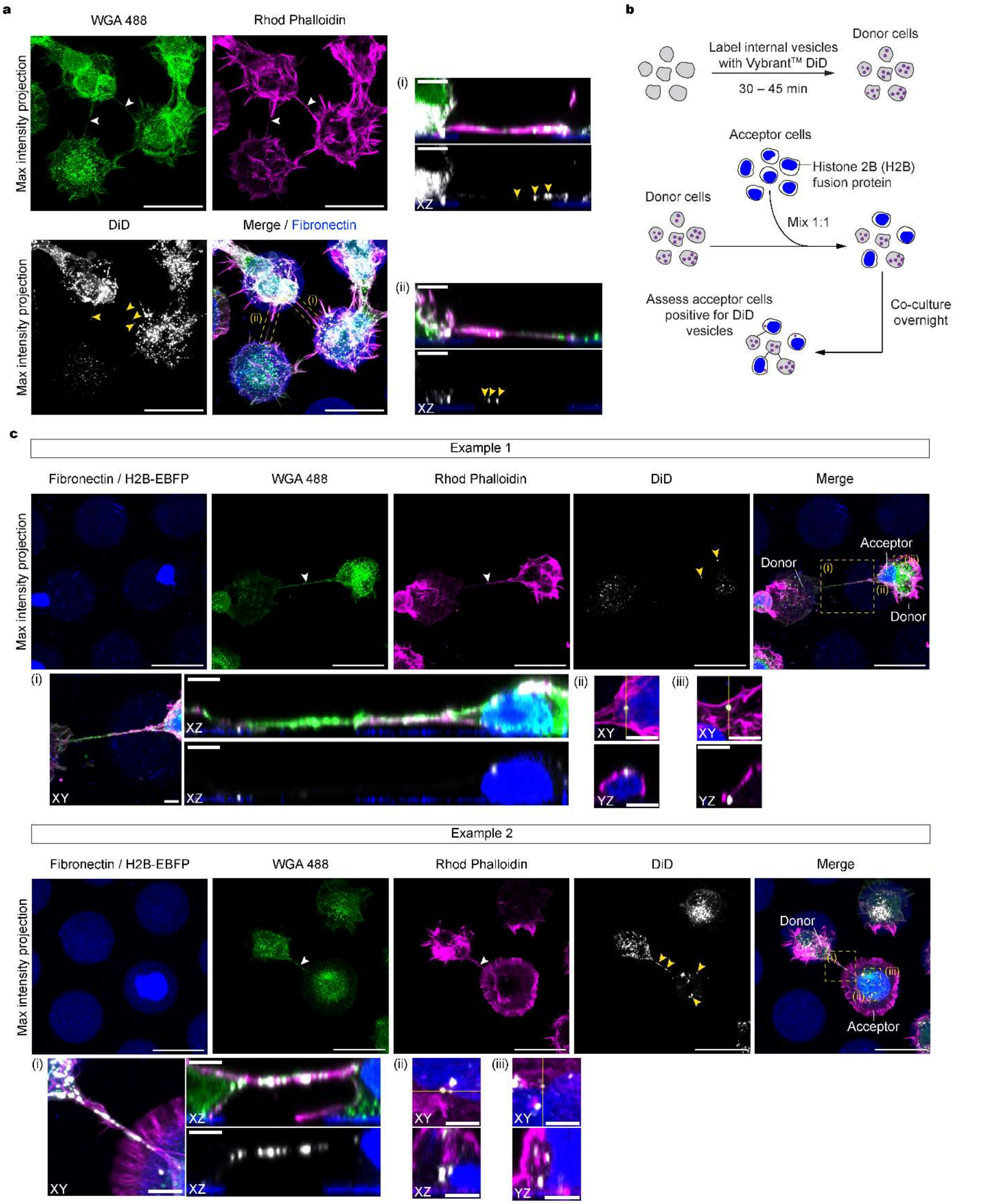


**Supplementary Fig. 2: Functional TNTs connecting micropatterned cells permit vesicle transfer.**

(**a**) Representative max intensity projections of TNTs connecting neighbouring CAD cells cultured on *D*15 micropatterns (fibronectin labelled with Alexa Flour 405) that contain DiD-labelled vesicles within them. Subpanels (i, ii) show the XZ projections through the axis of the TNTs indicated in the dashed yellow boxes. (**b**) Scheme depicting the co-culture experiment for assessing TNT functionality through DiD-labelled vesicle transfer between donor cells to acceptor cells expressing a fluorescently tagged histone 2b (H2B) fusion protein. (**c**) Representative examples of co-cultured CAD cells plated on *D*15 micropatterns (fibronectin labelled with Alexa Flour 405). Example 1 highlights a TNT linking a donor cell with an acceptor cell positive for several donor-derived vesicles. Example 2 shows a vesicle-positive TNT connecting a DiD-labelled donor cell and a H2B-EBFP expressing acceptor cell having received donor-derived vesicles. Dashed yellow boxes correspond to labelled subpanels (i−iii) showcasing zoom-ins and orthogonal projections of TNTs and DiD vesicles within acceptor cells; XZ projections of TNTs were made through the axis of the connection. TNTs are annotated with white arrowheads; yellow arrowheads point to DiD-labelled vesicles. Scale bars, 30 µm; Insets, 5 µm.

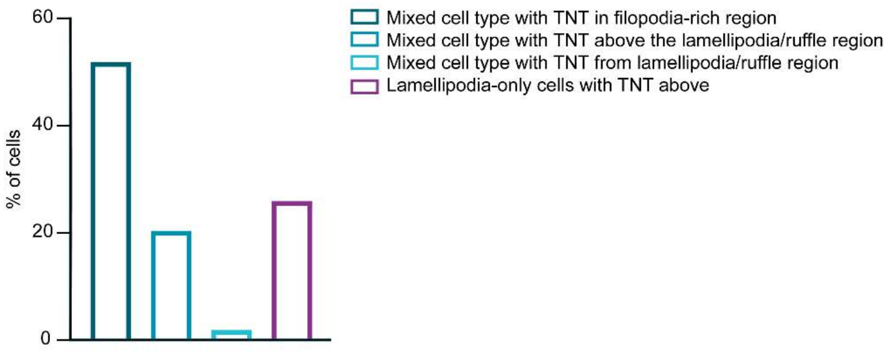


**Supplementary Fig. 3: Single-cell analysis of TNT origin.**
Bar graph showing the categorisation of TNTs and the cellular vicinity from which the TNT was originating. Single-cell analysis was performed on those cells classified in Fig. 2a,b with mixed (hairy and lamellipodial/ruffled) and lamellipodial-only phenotypes; TNTs were categorised as emanating from a filopodia-rich region, directly from a lamellipodia/ruffle region, or spatially separated above a lamellipodia/ruffle region.

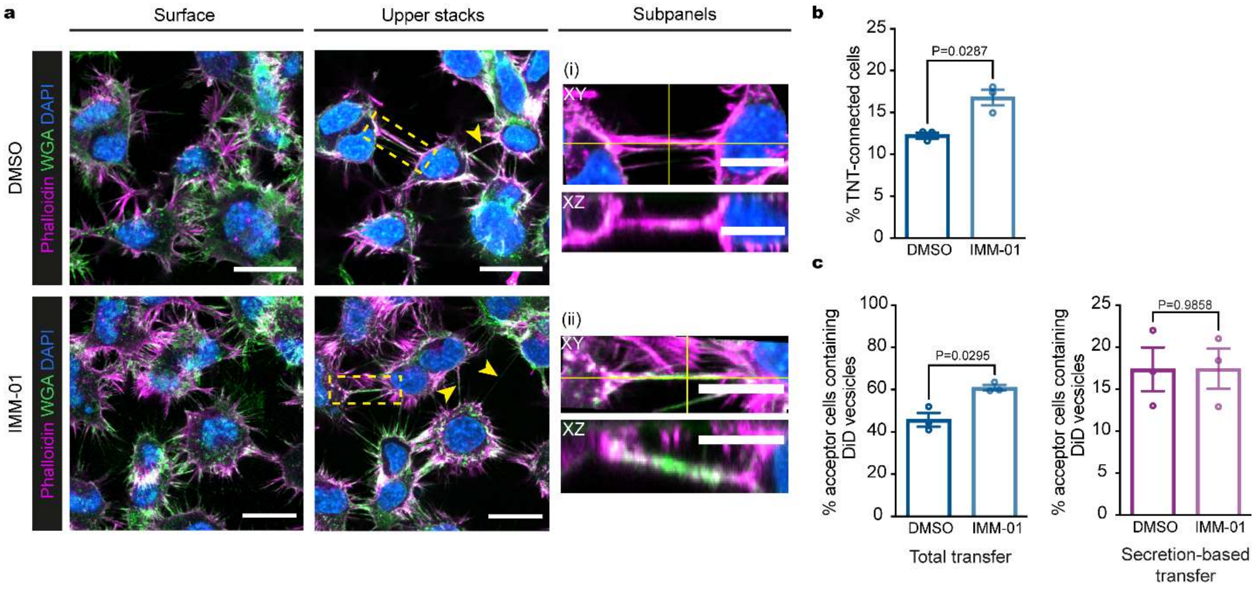


**Supplementary Fig. 4: Linear F-actin promotion leads to TNT-like structure formation.** **(a)** Representative images of surface and upper stacks of DMSO- and 1 µM IMM-01-treated cells plated on non-patterned surfaces. Yellow arrowheads annotate TNT-like protrusions. Subpanels (i, ii) show the XY and XZ projections through the axis of the TNTs indicated in the dashed yellow boxes. **(b)** Bar graph showing the quantification of TNT-like protrusions in DMSO (785 cells analysed; 12.3 ± 0.3%) and IMM-01 conditions (801 cells analysed; 16.8 ± 0.9%). Data are from 3 individual experiments and are represented as a mean ± SEM. Statistical analysis was performed using a t-test with Welch's correction, P = 0.0287. **(c)** Left: Bar graph showing the quantification of acceptor cells containing DiD-stained vesicles in the 1 µM IMM-01 treated co-culture. Data are from 3 individual experiments and are represented as a mean ± SEM. Statistical analysis was performed using a t-test with Welch's correction to compare IMM-01 (201 acceptor cells analysed; 61.0 ± 1.3%) with DMSO total co-culture transfer (202 acceptor cells analysed; 45.8 ± 3.3%), P = 0.0295. Right: Bar graph showing the quantification of acceptor cells containing DiD-stained vesicles obtained through secretion-based transfer (i.e., acceptor cells were cultured in conditioned media from DMSO- or IMM-01-treated donor cells). Data are from 3 individual experiments and are represented as a mean ± SEM. Statistical analysis was performed using a t-test with Welch's correction to compare DMSO (360 cells analysed; 17.4 ± 2.6%) versus IMM-01 (265 cells analysed; 17.4 ± 2.4%) conditions, P = 0.9858. Scale bars, 20 µm; Subpanels, 10 µm.

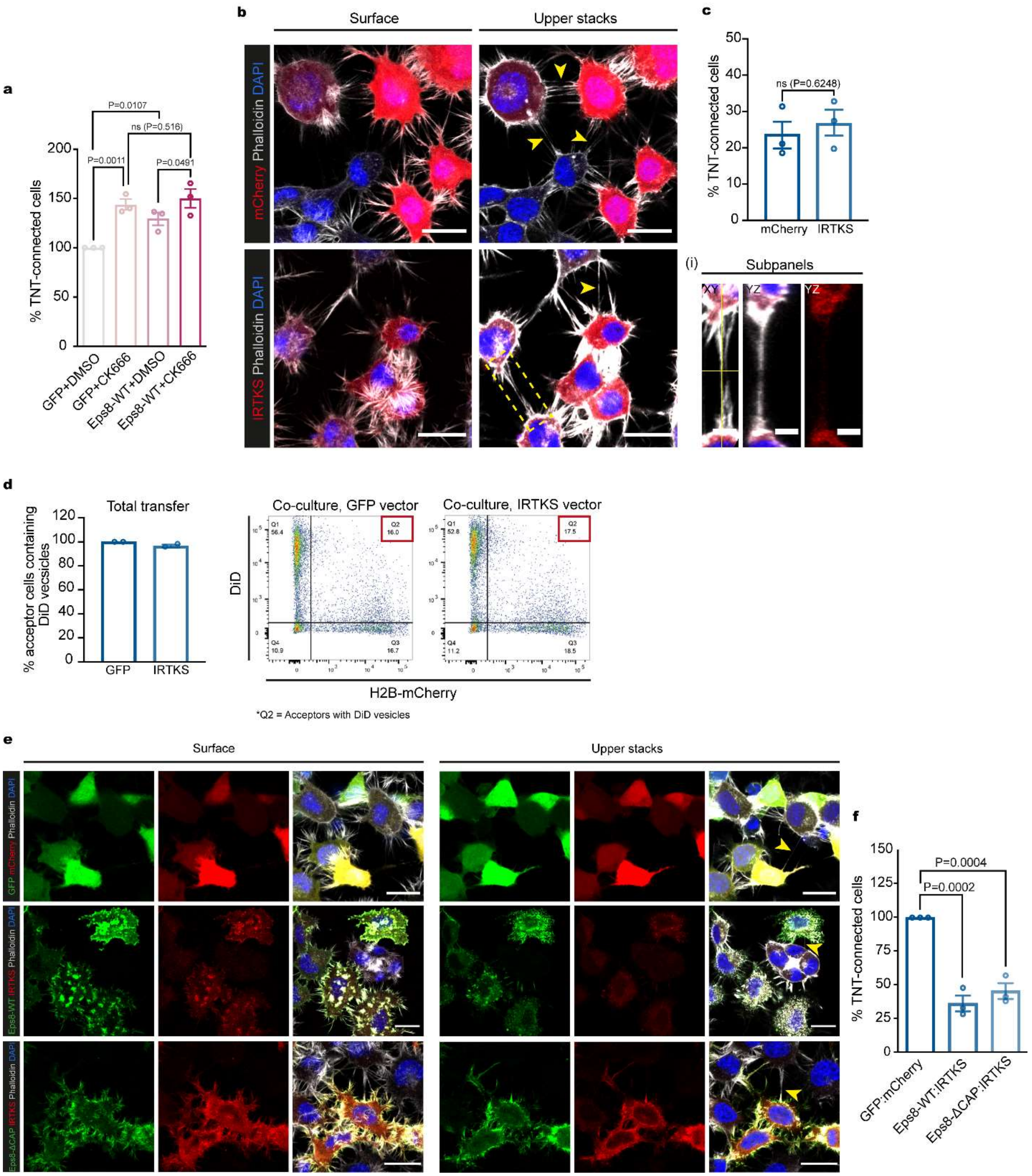


**Supplementary Fig. 5: IRTKS does not promote TNT formation, but rather decreases it when co-expressed with Eps8.**

**(a)** Bar graph showing the quantification of TNT-connected cells prior to immunoprecipitation for GFP + DMSO (297 cells analysed; 100%), GFP + CK-666 (229 cells analysed; 144.0 ± 5.4%), GFP-Eps8-WT + DMSO (224 cells analysed; 129.4 ± 6.2%) and GFP-Eps8-WT + CK-666 cells (246 cells analysed; 150.0 ± 9.5%). Data are from 3 individual experiments and are represented as a mean ± SEM. Statistical analysis was performed using an ordinary ANOVA with Tukey's multiple comparison test. P values for each comparison are stated on the bar graph. **(b)** Representative images of surface and upper stacks of mCherry-transfected control cells and IRTKS-mCherry transfected cells plated on non-patterned surfaces. Yellow arrowheads annotate TNT-like protrusions. Subpanel (i) shows the XY and YZ projections through the axis of the TNT indicated in the dashed yellow box. **(c)** Bar graph showing quantification of TNT-connected cells in mCherry control (309 cells analysed; 23.6 ± 4%) and in IRTKS-mCherry cells (268 cells analysed; 26.5 ± 3.9%). Data are from 3 individual experiments and are represented as a mean ± SEM. Statistical analysis was performed using a t-test with Welch's correction, P = 0.6248. **(d)** Left: Bar graph showing total transfer analysis in GFP control co-culture (100%) and IRTKS-GFP co-culture (96.5 ± 1.3%). Data are from 2 individual experiments and are represented as a mean ± SEM. Right: Gating strategy for flow cytometry measurements of total transfer. Q2 represents H2B-mCherry-labeled acceptor cells containing donor-derived DiD vesicles. **(e)** Left: Representative surface images of GFP:mCherry (control), GFP-Eps8-WT:IRTKS-mCherry and GFP-Eps8-ΔCAP:IRTKS-mCherry co-transfected cells. Right: Representative images of upper stacks of GFP:mCherry, GFP-Eps8-WT:IRTKS-mCherry and GFP-Eps8-ΔCAP:IRTKS-mCherry co-transfected cells. Cells were plated on non-patterned surfaces. **(f)** Bar graph showing the quantification of TNT-connected cells in GFP:mCherry (453 cells analyzed; 100%), Eps8-WT:IRTKS (203 cells analyzed; 36.0 ± 5.9%) and in Eps8-ΔCAP:IRTKS (252 cells analyzed; 45.1 ± 5.9%) co-transfected control cells. Data are from 3 individual experiments and are represented as a mean ± SEM. Statistical analysis was performed using an ordinary ANOVA with Dunnett's multiple comparison test. P values for each comparison are stated on the bar graph. Scale bars, 20 μm; Subpanels, 5 μm.

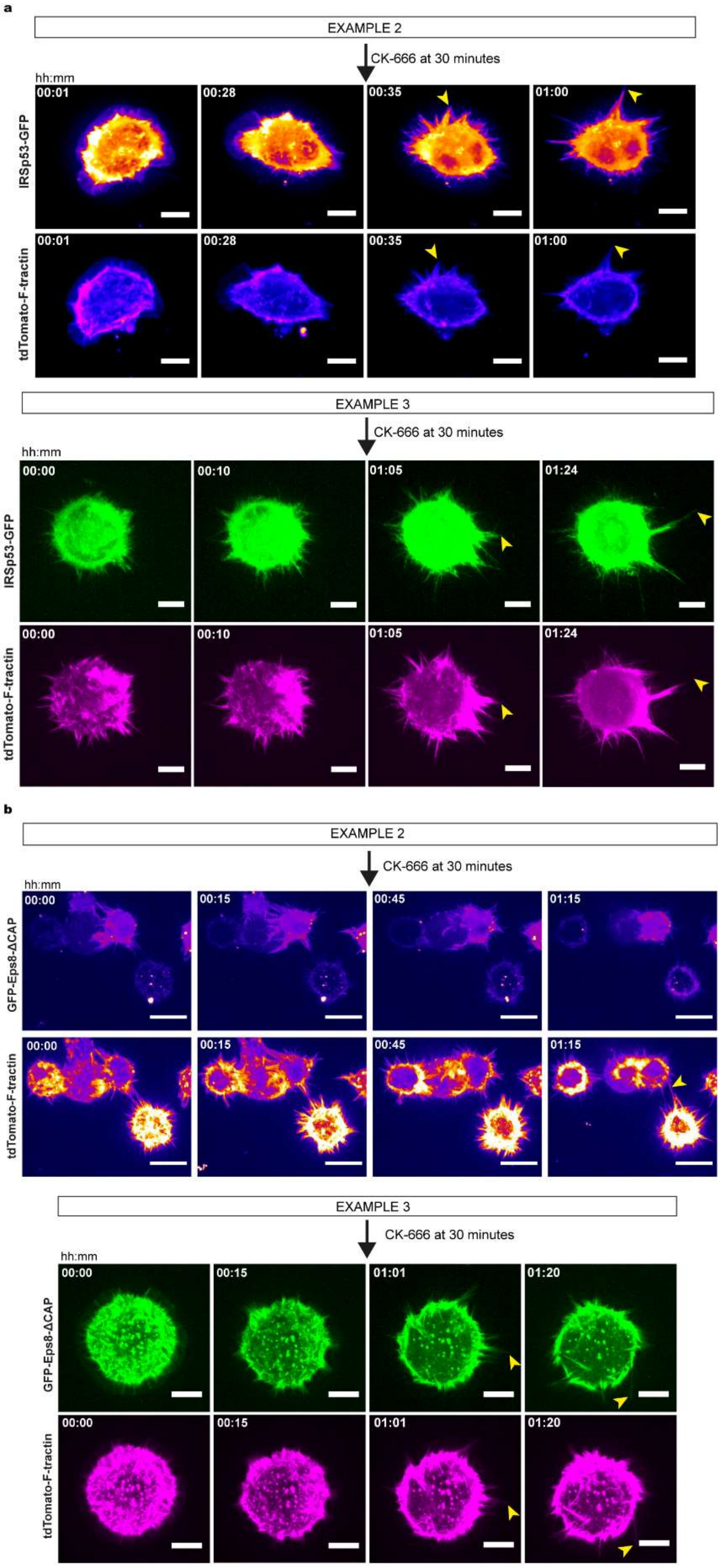
a
EXAMPLE 2
CK-666 at 30 minutes
hh:mm
00:01
00:28
00:35
01:00
IRSp53-GFP
tdTomato-F-tractin
EXAMPLE 3
CK-666 at 30 minutes
hh:mm
00:00
00:10
01:05
01:24
IRSp53-GFP
tdTomato-F-tractin
b
EXAMPLE 2
CK-666 at 30 minutes
hh:mm
00:00
00:15
00:45
01:15
GFP-Eps8-ΔCAP
tdTomato-F-tractin
EXAMPLE 3
CK-666 at 30 minutes
hh:mm
00:00
00:15
01:01
01:20
GFP-Eps8-ΔCAP
tdTomato-F-tractin

**Supplementary Fig. 6: Eps8 and IRSp53 are recruited to form longer protrusions upon Arp2/3 inhibition.**

(**a**) Representative time-lapse images of protrusion formation in IRSp53-transfected cells before and after the addition of CK-666 (at 30 min). Example 2 and Example 3 corresponds to Supplementary Video 9 and Supplementary Video 10, respectively. **(b)** Representative time-lapse images of protrusion formation in Eps8-ΔCAP-transfected cells before and after the addition of CK-666 (at 30 min). Example 2 and Example 3 corresponds to Supplementary Video 11 and Supplementary Video 12, respectively. Both IRSp53- and Eps8-ΔCAP-transfected cells additionally expressed the F-actin label tdTomato-F-tractin. Yellow arrowheads throughout show protrusions that are formed after CK-666 addition. Scale bars, 10 µm.

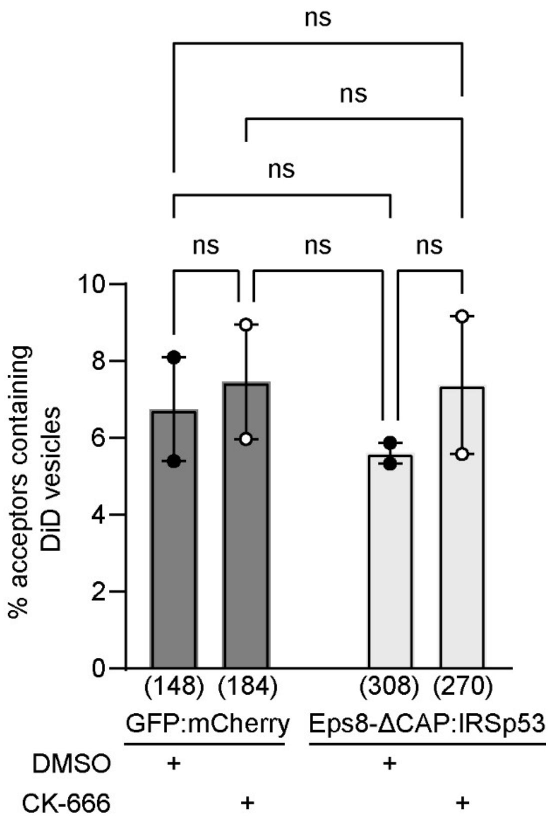


**Supplementary Fig. 7: Secretion-based transfer of DiD-labelled vesicles is invariant to CK-666 treatment.**

Plot of the percentage of acceptor cells containing DiD-labelled vesicles from secretion-based transfer experiments for GFP:mCherry + DMSO (6.8 ± 1.4%), GFP:mCherry + CK-666 (7.5 ± 1.5%), Eps8-ΔCAP:IRSp53 + DMSO (5.6 ± 0.3%), and Eps8-ΔCAP:IRSp53 + CK-666 (7.4 ± 1.8%) (mean ± SEM). Data was from two individual experiments and the total number of acceptor cells analysed in each condition is indicated below. Statistical analysis was performed using a Kruskal-Wallis multiple comparison test. P value for all comparisons > 0.9999 except GFP:mCherry + CK-666 vs. Eps8-ΔCAP:IRSp53 + DMSO was P = 0.9183; ns = non-significant.

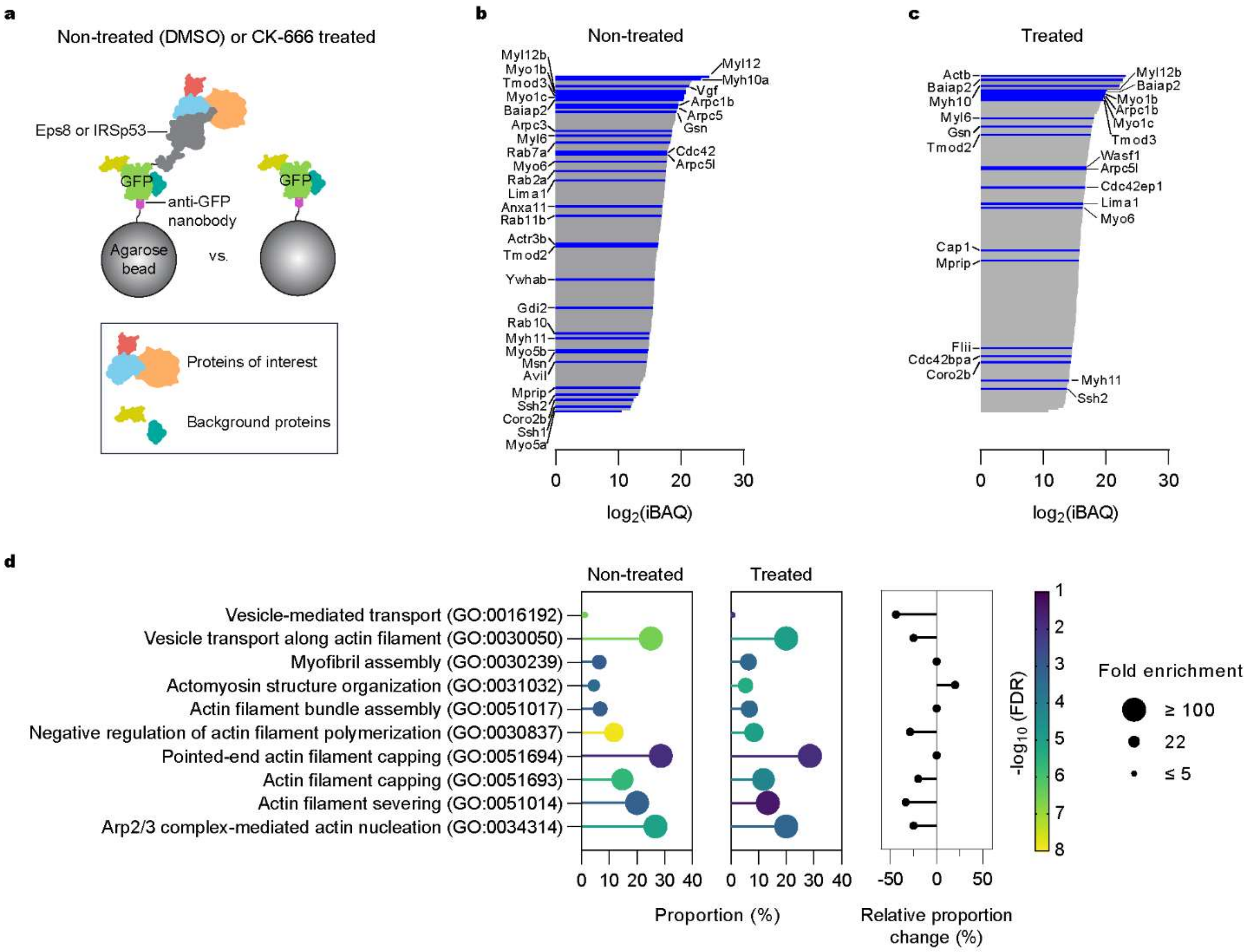


**e**

| GO Biological Process | Ref Count | Proteins |
|---|---|---|
| Vesicle-mediated transport (GO:0016192) | 1383 | Actb, Anxa1, Cdc42, Gsn, Cap1, Myo1b, Myo5a, Myo5b, Myo6 Myh9, Myh10, Gdi2, Rab2a, Rab7a, Rab10, Rab11b, Wasf1 |
| Vesicle transport along actin filament (GO:0030050) | 20 | Myo1b, Myo1c, Myo5a, Myo5b, Myo6 |
| Myofibril assembly (GO:0030239) | 63 | Flii, Tmod2, Tmod3, Myh10 |
| Actomyosin structure organization (GO:0031032) | 113 | Flii, Tmod2, Tmod3, Myh9, Myh10, Cdc42bpa |
| Actin filament bundle assembly (GO:0051017) | 60 | Myh10, Lima1, Myo1b, Baiap2 |
| Negative regulation of actin filament polymerization (GO:0030837) | 60 | Ssh1, Ssh2, Flii, Gsn, Avil, Tmod2, Tmod3 |
| Pointed-end actin filament capping (GO:0051694) | 7 | Tmod2, Tmod3 |
| Actin filament capping (GO:0051693) | 34 | Flii, Gsn, Avil, Tmod2, Tmod3 |
| Actin filament severing (GO:0051014) | 15 | Flii, Gsn, Avil |
| Arp2/3 complex-mediated actin nucleation (GO:0034314) | 15 | Arpc1b, Arpc3, Arpc5, Arpc5l |

■ Non-treated only ■ Treated only ■ Common

**Supplementary Fig. 8: Eps8-related proteomic data relevant for Figure 6.**
(**a**) Schematic depicting the GFP-Trap immunoprecipitation strategy for identifying actin-related protein hits associated with Eps8 and IRSp53 "bait" proteins. (**b**) Actin-related proteins identified as differentially abundant in Eps8-WT as compared to the negative GFP control for non-treated (DMSO) cells. (**c**) Actin-related proteins identified as differentially abundant in Eps8-WT as compared to the negative GFP control for CK-666-treated cells. (**b**, **c**) Plots of the intensity-based absolute quantification (iBAQ) values were generated from the sum of peak intensities of all peptides corresponding to a specific protein divided by the number of observable peptides. (**d**, **e**) Results of a gene ontology (GO) biological process term analysis for non-treated and CK-666-

treated Eps8 samples. (**d**) Left: Proportion of protein hits for a given GO term. The number of protein hits in the network for a given GO term was normalized by the total number of proteins assigned to a given GO term using the whole mouse genome as a reference (Ref count in **e**). The graph is coloured by the false discovery rate (FDR). Sizes reflect the fold enrichment of the number proteins in the network divided by the number of proteins expected to be annotated with a given GO term in a randomly generated network of the same size. Right: Relative change in the proportion of proteins in a GO term when comparing CK-666-treated to non-treated Eps8-WT expressing CAD cells. (**e**) Table summarizing mapped proteins to their corresponding GO term for **d**. The number of reference proteins in the mouse genome for a given GO term is provided (Ref count). Colour code: Teal, proteins only present in the non-treated (DMSO) Eps8 pull down; Magenta, proteins only present in the CK-666-treated Eps8 pull down; Black, proteins common to both pull-downs.

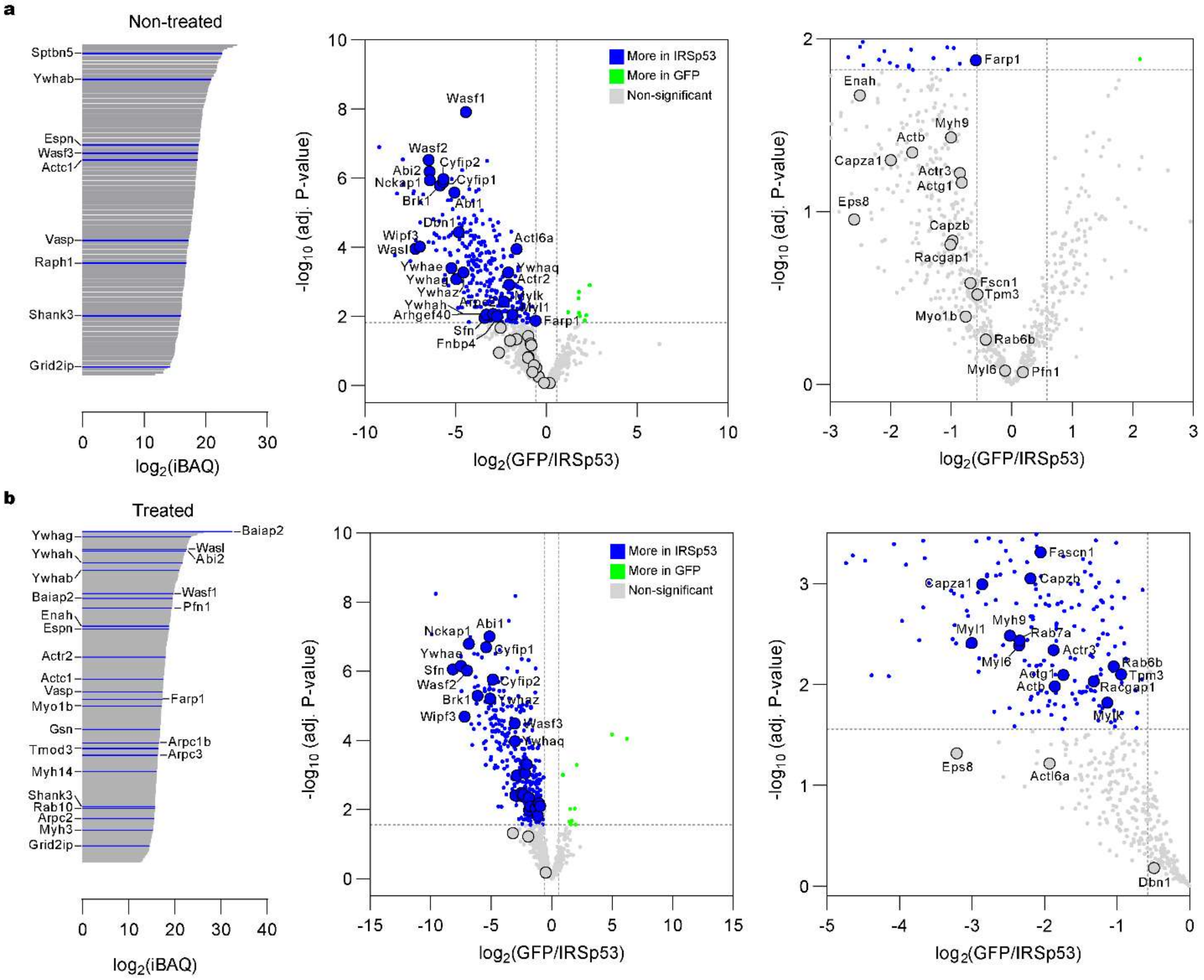


**Supplementary Fig. 9: IRSp53-related proteomic data relevant for Figure 7.**
Actin-related proteins identified in IRSp53 as compared to the negative GFP control for non-treated (DMSO) (**a**) and CK-666-trreated (**b**) CAD cells. iBAQ plots present proteins only found to be present in their respective IRSp53 samples as compared to the negative GFP control. Volcano plots show the differential abundance of identified proteins that are more present in respective IRSp53 samples.

**Supplementary Video 1.**
Time-lapse imaging showing CAD cell adherence to fibronectin micropatterns false-coloured in blue (rhodamine-labelled fibronectin). Merged brightfield and fluorescence channels are presented on the left and the RICM channel is presented on the right. Frames were acquired at a 1-min interval over 5 hr and corrected post acquisition for lateral drift. Video playback speed is at 7 fps. Time in hh:mm. Scale bar, 10 µm. See selected still frames in Supplementary Fig. 1d.

**Supplementary Video 2.**
Tracking of CAD cell movement on fibronectin micropatterns using video microscopy. Video depicts a representative example of a well-confined CAD cell. Merged brightfield and fluorescence channels are presented with the micropatterns false-coloured in blue (rhodamine-labelled fibronectin). The reference fluorescence image of the micropatterns was taken at the first timepoint only. The cell's tracked centre-of-mass is overlaid as a rainbow-coloured trajectory. Frames were acquired at a 5-min interval over 18 hr. Video playback speed is at 12 fps. Scale bar, 10 µm. See selected still frames and a plot of the trajectory in Supplementary Fig. 1g, Example 1.

**Supplementary Video 3.**
Tracking of CAD cell movement on fibronectin micropatterns using video microscopy. Video shows a representative example that cells remain confined even after division. Merged brightfield and fluorescence channels are presented with the micropatterns false-coloured in blue (rhodamine-labelled fibronectin). The reference fluorescence image of the micropatterns was taken at the first timepoint only. The cell's tracked centre-of-mass is overlaid as a rainbow-coloured trajectory. Frames were acquired at a 5-min interval over 18 hr. Video playback speed is at 12 fps. Scale bar, 10 µm. See selected still frames and a plot of the trajectory in Supplementary Fig. 1g, Example 2.

**Supplementary Video 4.**
Tracking of CAD cell movement on fibronectin micropatterns using video microscopy. Video shows a rare example of a CAD cell moving between adjacent micropatterns. Merged brightfield and fluorescence channels are presented with the micropatterns false-coloured in blue (rhodamine-labelled fibronectin). The reference fluorescence image of the micropatterns was taken at the first timepoint only. The cell's tracked centre-of-mass is overlaid as a rainbow-coloured trajectory. Frames were acquired at a 5-min interval over 18 hr. Video playback speed is at 12 fps. Scale bar, 10 µm. See selected still frames and a plot of the trajectory in Supplementary Fig. 1g, Example 3.

**Supplementary Video 5.**
Time-lapse imaging of a mock-treated (DMSO) CAD cell expressing EGFP F-Tractin (green). Cell Mask$^{TM}$ Deep Red (magenta) was used to visualize the plasma membrane and the pulled nanotube. Frames were acquired at an interval of 30 sec and video playback speed is at 3 fps. Time in hh:mm:ss. Scale bar, 5 µm.

**Supplementary Video 6.**
Time-lapse imaging of a CAD cell expressing EGFP F-Tractin (green) treated with 50 µM CK-666. Cell Mask$^{TM}$ Deep Red (magenta) was used to visualize the plasma membrane and the pulled nanotube. Frames were acquired at an interval of 30 sec and video playback speed is at 3 fps. Time in hh:mm:ss. Scale bar, 5 µm.

**Supplementary Video 7.**
Time-lapse imaging of a CAD cell expressing IRSp53-GFP 30 min prior to and upon treatment with 50 μM CK-666. Frames were acquired at an interval of 30 sec and video playback speed is at 7 fps. Time in hh:mm:ss. Scale bar, 10 μm.

**Supplementary Video 8.**
Time-lapse imaging of a CAD cell expressing GFP-Eps8-ΔCAP 30 min prior to and upon treatment with 50 μM CK-666. Frames were acquired at an interval of 1 min and video playback speed is at 7 fps. Time in hh:mm. Scale bar, 20 μm.

**Supplementary Video 9.**
Time-lapse imaging of a CAD cell expressing IRSp53-GFP (left) and tdTomato-F-tractin (right) prior to and upon treatment with 50 μM CK-666. Cells were sparsely plated on non-micropatterned surfaces. Frames were acquired at an interval of 1 min and video playback speed is at 7 fps. Time in hh:mm. Scale bar, 10 μm.

**Supplementary Video 10.**
Time-lapse imaging of a CAD cell expressing IRSp53-GFP (left) and tdTomato-F-tractin (right) prior to and upon treatment with 50 μM CK-666 on *D*40 micropatterns (not visible). Frames were acquired at an interval of 1 min and video playback speed is at 7 fps. Time in hh:mm. Scale bar, 10 μm.

**Supplementary Video 11.**
Time-lapse imaging of a CAD cell expressing GFP-Eps8-ΔCAP (left) and tdTomato-F-tractin (right) 30 min prior to and upon treatment with 50 μM CK-666. Cells were sparsely plated on non-micropatterned surfaces. Frames were acquired at an interval of 1 min and video playback speed is at 7 fps. Time in hh:mm. Scale bar, 10 μm.

**Supplementary Video 12.**
Time-lapse imaging of a CAD cell expressing GFP-Eps8-ΔCAP (left) and tdTomato-F-tractin (right) 30 min prior to and upon treatment with 50 μM CK-666 on *D*40 micropatterns (not visible). Frames were acquired at an interval of 1 min and video playback speed is at 7 fps. Time in hh:mm. Scale bar, 10 μm.
